# The ammonia–senescence ratchet drives an adaptation–senescence– ammonioptosis transition in liver cancer

**DOI:** 10.64898/2026.10.11.756515

**Authors:** Yubin Han, Miao Huang, Yulin Xie, Shanshan Zhang, Jinshen Wang, Lin Han, Chunxia Li, Yong Zhang

**Affiliations:** Institute of Frontier Chemistry, School of Chemistry and Chemical Engineering, Shandong University, Qingdao 266237, P. R. China; Institute of Marine Science and Technology, Shandong University, Qingdao 266237, P. R. China; Shandong Institutes of Industrial Technology, Jinan 250101, P. R. China; Department of Gastrointestinal Surgery, Shandong Provincial Hospital Affiliated to Shandong First Medical University, Jinan 250021, P. R. China; Department of Biomedical Engineering, College of Biomedicine, The City University of Hong Kong, Kowloon 999077, Hong Kong

## Abstract

Senescence suppresses tumor proliferation while exposing survival dependencies, motivating separate, sequential steps of senescence induction and senescent-cell clearance. Whether senescence-sustaining metabolism can instead be overdriven to trigger cell-autonomous death remains unknown. Senescent cells generate ammonia through glutaminolysis to buffer intracellular acidification, yet extracellular ammonia can impair anti-tumor immunity, leaving unexplored whether ammonia can instead drive tumor-cell-intrinsic death. Here, we reverse this logic and identify a self-reinforcing ammonia–senescence ratchet in which ammonia accumulation promotes senescence, whereas senescence reciprocally enhances ammoniagenesis, progressively driving tumor cells from metabolic adaptation through senescence to ammonioptosis. Reducing intracellular ammonia suppressed senescence and cell death, whereas ammonia repletion restored lethality. We engineered acid-responsive H3B-120@CoGd-LDH (HBCG) to preferentially amplify tumor-cell ammonia by reinforcing senescence and ammoniagenesis, restricting ammonia disposal and weakening lysosomal buffering. HBCG limited extracellular ammonia accumulation and CD8⁺ T-cell exposure in co-cultures, induced ammonioptosis in patient-derived liver-cancer organoids and suppressed tumor growth in immunocompetent mice. This work reveals that adaptation, senescence and ammonioptosis can be connected through escalating intracellular ammonia stress, providing a mechanistic framework for understanding stress-driven tumor-cell fate transitions.

---

Cellular senescence durably arrests tumor cells while exposing state-specific metabolic dependencies, but leaves persistent senescent populations that often require subsequent clearance^1, 2^. This limitation has motivated “one–two-punch” strategies, in which senescence induction is followed by a mechanistically distinct intervention that exploits vulnerabilities acquired during senescence^3–8^. Yet these approaches treat senescence induction and senescent-cell elimination as separate therapeutic steps. Whether senescence-sustaining programs can instead be overdriven to make senescence self-terminating remains unknown.

Senescent cells undergo extensive metabolic remodeling, including enhanced glutaminase 1 (GLS1)-dependent glutaminolysis, which continuously generates ammonia to counteract intracellular acidification and support survival^9^. Yet the biological consequences of ammonia are highly context dependent. Within tumors, ammonia can be reutilized to support biosynthesis and growth, whereas its accumulation in the microenvironment impairs effector T-cell function and favours regulatory T-cell adaptation^10–12^. Accordingly, therapeutic strategies have largely focused on ammonia depletion or detoxification, leaving unresolved whether tumor-derived ammonia can instead be redirected from a diffusible immunosuppressive liability into a tumor-cell-intrinsic lethal stress.

Here we show that intracellular ammonia and senescence form a self-reinforcing metabolic ratchet that propels tumor cells from metabolic adaptation through senescence towards ammonioptosis. Restricting carbamoyl phosphate synthetase 1 (CPS1)-dependent ureagenic disposal causes intracellular ammonia accumulation and promotes senescence, whereas senescent tumor cells reciprocally increase GLS1-dependent ammoniagenesis, further escalating intracellular ammonia to trigger ammonioptosis, a senescence-coupled, ammonia-dependent tumor-cell death (Fig. 1). Lowering intracellular ammonia suppressed both senescence and subsequent cell death, whereas ammonia repletion restored the lethal phenotype. Inhibitors of apoptosis, ferroptosis or necroptosis provided little protection. Together, these findings establish intracellular ammonia as a determinant of tumor-cell fate and reveal that senescence-sustaining metabolism can be overdriven to precipitate cell-autonomous death.

**Fig. 1.**
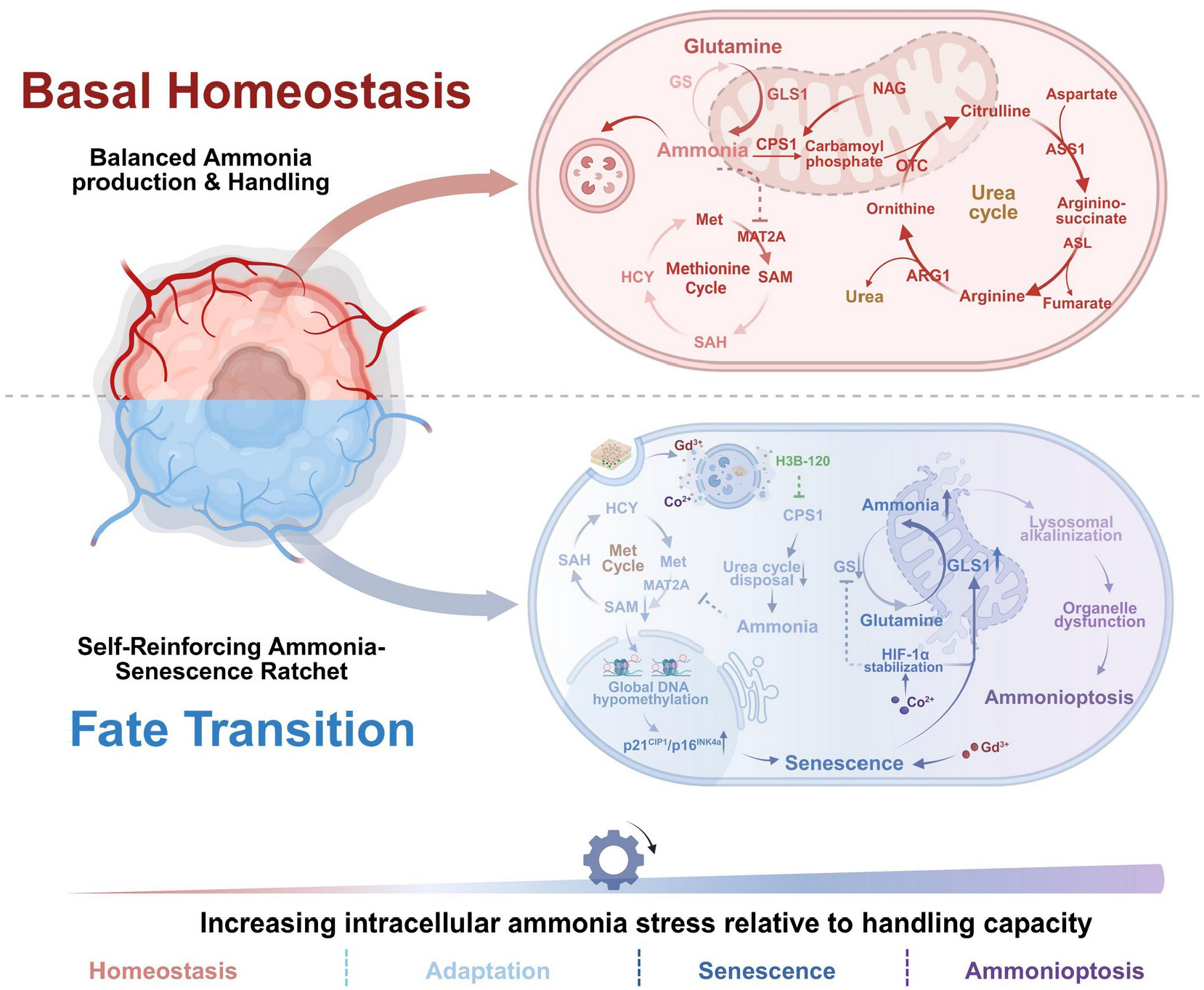
The ammonia–senescence ratchet drives an adaptation–senescence–ammonioptosis transition in liver cancer. Scheme of intracellular ammonia crossing successive cell-state thresholds. Under basal homeostasis, GLS1-dependent glutaminolysis is balanced by CPS1-dependent ureagenesis, glutamine synthetase (GS)-mediated reassimilation and compartmental buffering, thereby maintaining methionine adenosyltransferase 2A (MAT2A)-dependent S-adenosylmethionine (SAM) production and global DNA methylation; Following HBCG uptake, acidic conditions trigger H3B-120 release, inhibiting CPS1-dependent ureagenesis and driving intracellular ammonia accumulation, which suppresses the methionine adenosyltransferase 2A (MAT2A)–S-adenosylmethionine (SAM) axis, leading to DNA hypomethylation and p16^INK4a^/p21^CIP1^-associated senescence. Senescent cells further increase GLS1-dependent ammoniagenesis, forming a self-reinforcing ammonia–senescence ratchet. HBCG amplifies this feedback through Co^2+^-associated HIF-1α stabilization and GLS1 induction, while Gd^3+^ reinforces senescence and the layered double hydroxide (LDH) component disrupts lysosomal buffering. Continued ammonia accumulation beyond lysosomal buffering capacity causes lysosomal dysfunction and mitochondrial injury, culminating in ammonioptosis. The bottom gradient denotes increasing intracellular ammonia stress.

To exploit this spatially divergent biology, we engineered acid-responsive H3B-120@CoGd layered double hydroxide (HBCG) to preferentially amplify tumor-cell-intrinsic ammonia while limiting immune-cell exposure. Unlike urease-based systems that generate ammonia from extracellular substrates, HBCG acts on tumor-intrinsic nitrogen metabolism by simultaneously reinforcing senescence and ammoniagenesis, restricting ammonia disposal and weakening lysosomal buffering. In tumor–T-cell co-cultures, HBCG produced a tumor-cell-biased ammonia distribution, with pronounced ammonia accumulation in tumor cells but limited increases in the shared medium and CD8⁺ T cells. HBCG induced ammonioptosis in Hepa1-6 cells, patient-derived hepatocellular carcinoma and intrahepatic cholangiocarcinoma organoids and immunocompetent tumors. Thus, spatially biased amplification converts ammonia from a senescence-supporting metabolite and extracellular immune liability into a tumor-cell-intrinsic lethal stress.

## CPS1 blockade restricts ureagenic ammonia disposal and promotes ammonia retention beyond adaptive capacity

Glutamine-addicted liver cancer cells continuously generate ammonia through glutaminolysis, yet this stress remains compatible with an adaptive state while metabolic disposal, reassimilation and buffering match production. In hepatocytes, mitochondrial entry into the urea cycle provides a principal route for high-capacity ammonia disposal. CPS1 catalyzes the first committed and rate-limiting step of ureagenesis, incorporating ammonia and bicarbonate into carbamoyl phosphate at the expense of two ATP molecules^13^. This mitochondrial reaction represents a major entry point for excess nitrogen into the urea cycle and is therefore critical for maintaining intracellular and systemic ammonia homeostasis. Carbamoyl phosphate is subsequently processed through downstream urea-cycle reactions to convert nitrogen into urea for systemic excretion. CPS1 activity is tightly regulated by N-acetylglutamate (NAG), an obligate allosteric activator synthesized from glutamate and acetyl-CoA by N-acetylglutamate synthase (NAGS)^14, 15^. We therefore investigated whether pharmacological CPS1 inhibition would lower ammonia-handling capacity sufficiently for ongoing ammoniagenesis to exceed the adaptive range (Fig. 2a).

**Fig. 2.**
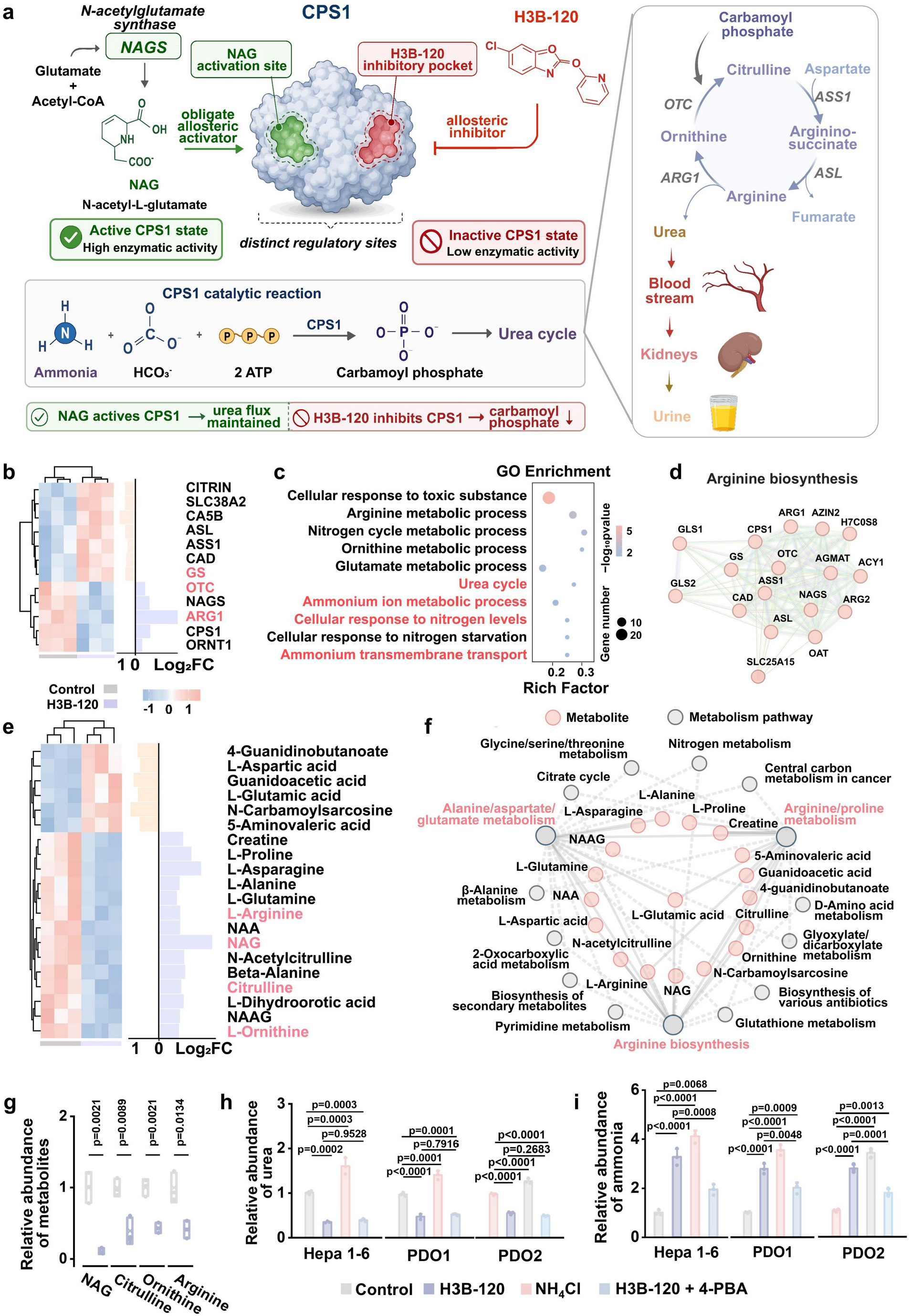
CPS1 blockade restricts ureagenic ammonia disposal and promotes intracellular ammonia accumulation. **a**, Schematic showing the allosteric regulation of CPS1 by the obligate activator N-acetyl-L-glutamate (NAG) and pharmacological inhibition by H3B-120 at a distinct inhibitory pocket. CPS1 catalyzes the ATP-dependent conversion of ammonia and bicarbonate into carbamoyl phosphate, initiating urea-cycle nitrogen disposal. **b**, Hierarchical clustering heat map of selected urea-cycle and nitrogen-handling genes in PDO1 following control or H3B-120 treatment, with corresponding log_2_ fold changes shown on the right. **c**, Gene Ontology (GO) enrichment analysis of differentially expressed genes identified by RNA sequencing in **b**. Dot size indicates gene number, colour denotes −log_10_(*P* value), and rich factor represents the proportion of differentially expressed genes assigned to each term. **d**, STRING protein–protein interaction network of arginine-biosynthesis-associated genes altered following H3B-120 treatment in PDO1. **e**, Heat map of targeted intracellular metabolites altered by H3B-120 treatment, highlighting urea-cycle intermediates and related amino-acid metabolites; corresponding log_2_ fold changes are shown on the right. **f**, Integrated metabolite–pathway network linking differentially abundant metabolites to enriched metabolic pathways following CPS1 blockade. Pink nodes indicate metabolites and grey nodes indicate metabolic pathways. **g**, Targeted metabolomic quantification of NAG, citrulline, ornithine and L-arginine in PDO1 following control or H3B-120 treatment. **h**, **i**, Relative intracellular urea (h) and ammonia (i) abundance in Hepa1-6 cells, PDO1 and PDO2 following treatment with control, H3B-120, NH_4_Cl or H3B-120 plus 4-PBA. Data in g–i are mean ± s.d.; individual points represent biologically independent samples, with *n* = 3 per condition. Statistical significance was assessed using two-sided unpaired Student’s *t*-tests in g and one-way ANOVA followed by Tukey’s multiple-comparisons test in h,i. Exact *P* values are shown in the corresponding panels.

PDOs were established from surgically resected liver tumor specimens (Supplementary Fig. 1). We treated Hepa 1-6 cells and PDOs with H3B-120, a selective CPS1 inhibitor that engages an allosteric pocket distinct from the NAG-binding site^16^. Parallel transcriptomic and metabolomic profiling was then performed in H3B-120-treated PDO1 using untreated organoids as controls. Differential expression analysis revealed coordinated suppression of the ureagenic program. Expression of NAGS and CPS1, which control mitochondrial urea-cycle entry, was reduced together with ARG1, which catalyzes the terminal production of urea. GS, which provides an alternative route for ammonia reassimilation, showed only a modest increase, consistent with engagement of a limited compensatory response (Fig. 2b). GO and KEGG analyses linked the transcriptional response to the urea cycle, ammonium metabolism and transport, and interconnected amino acid pathways, whereas PPI analysis organized differentially expressed arginine-biosynthesis genes into a connected network (Fig. 2c, d; Supplementary Fig. 2). Targeted metabolomics further revealed depletion of NAG and urea-cycle metabolites, including citrulline, ornithine and arginine, with integrated analysis mapping these changes to arginine biosynthesis and connected nitrogen-metabolic networks (Fig. 2e–g). Together, these transcriptional and metabolic changes indicated reduced ureagenic capacity, with NAG depletion imposing an additional constraint on residual CPS1 function.

To determine whether this molecular suppression shifted ammonia stress beyond adaptation, we measured intracellular ammonia and urea production. NH_4_Cl, used to impose exogenous ammonia stress, increased intracellular ammonia without reducing urea production in Hepa1-6 cells, PDO1 or PDO2, indicating that acute ammonia stress did not disable ureagenic disposal. By contrast, H3B-120 simultaneously reduced urea production and increased intracellular ammonia in all three models, indicating that ongoing ammoniagenesis had become uncoupled from ureagenic disposal (Fig. 2h, i). We next investigated whether lowering the ammonia burden could reverse this phenotype independently of CPS1. Treatment with 4-phenylbutyric acid (4-PBA) markedly reduced H3B-120-induced ammonia accumulation but did not restore urea abundance, indicating that ammonia could be relieved through an alternative nitrogen-disposal route while CPS1-dependent ureagenesis remained constrained. This separation between persistent suppression of urea production and partial normalization of intracellular ammonia further supports direct CPS1 inhibition as the primary defect caused by H3B-120. Together, these findings show that H3B-120 limits ureagenic ammonia disposal, allowing ongoing glutaminolytic ammoniagenesis to generate a sustained intracellular ammonia burden that exceeds the adaptive buffering capacity of liver tumor cells.

## Ammonia retention suppresses MAT2A–SAM methyl-donor axis and promotes senescence

Having shown that CPS1 inhibition shifted intracellular ammonia beyond adaptive handling capacity, we next examined how this metabolic stress altered downstream metabolism and cell state. We hypothesized that ammonia accumulation limits MAT2A-dependent SAM synthesis, thereby reducing methyl-donor availability and promoting senescence-associated cell-cycle arrest. Integrated transcriptomic, metabolomic and phenotypic analyses were performed. Transcriptomic profiling of H3B-120-treated PDO1 revealed suppression of one-carbon and DNA-methylation programs together with activation of DNA-damage, G1/S-checkpoint and senescence-associated programs (Fig. 3b). GO and KEGG enrichment analyses identified pathways associated with DNA replication, p53 and DNA-damage signaling, G1/S-transition control, one-carbon metabolism and SAM metabolism (Fig. 3c; Extended Data Fig. 1a). PPI network analysis further organized the differentially expressed genes into interconnected modules associated with methionine metabolism, DNA methylation, DNA replication and senescence-associated cell-cycle arrest (Fig. 3d). Among these changes, MAT2A was prominently downregulated. Because MAT2A catalyzes the ATP-dependent conversion of methionine into SAM^17^, its suppression provided a potential link between ammonia retention and cellular methyl-donor availability.

**Fig. 3.**
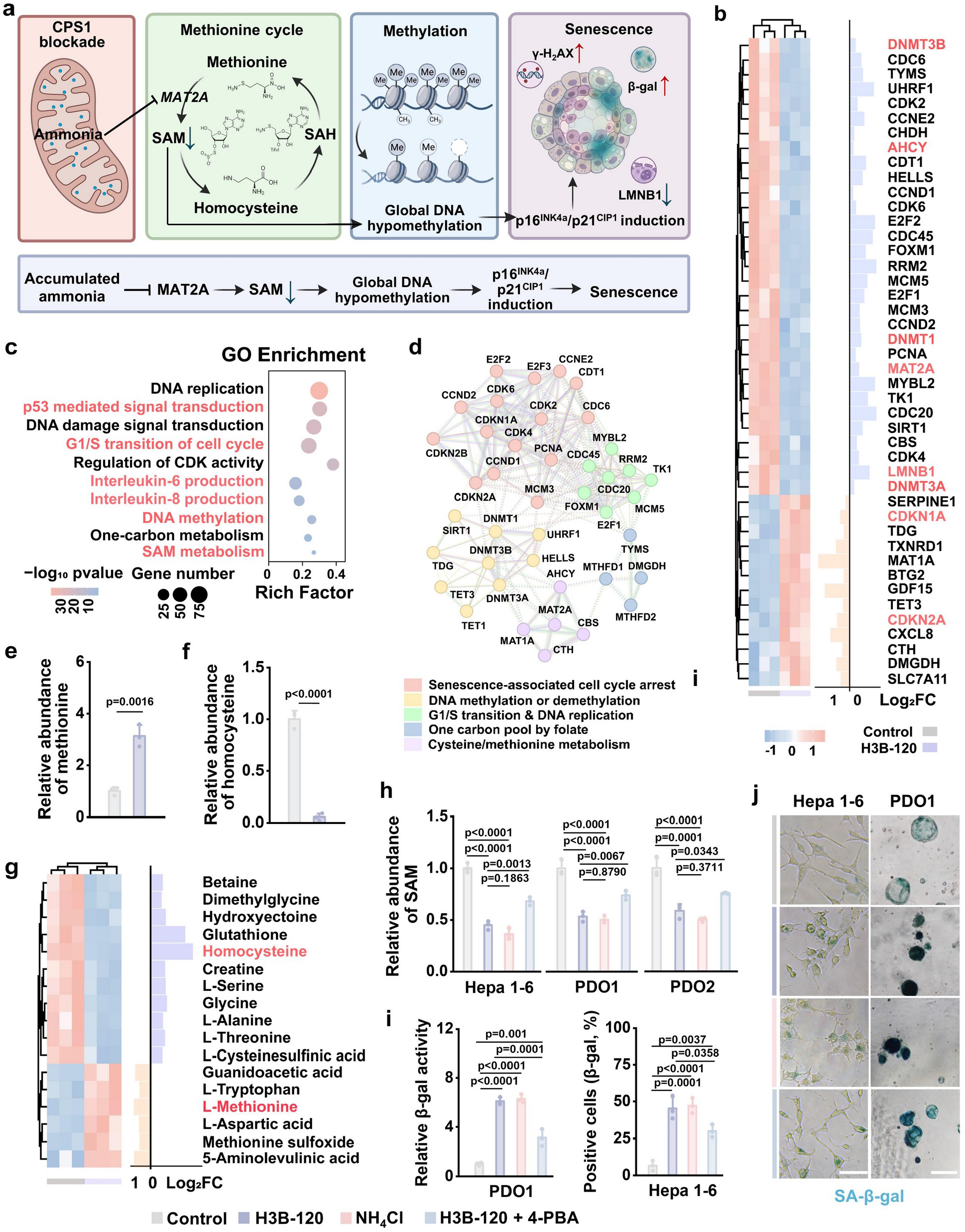
Ammonia accumulation suppresses the MAT2A–SAM methyl-donor axis and promotes senescence. **a**, Proposed model linking CPS1 blockade to ammonia retention, MAT2A suppression, reduced SAM availability, global DNA hypomethylation and senescence. **b**, Hierarchically clustered heat map of DNA-methylation, cell-cycle and senescence-related genes in H3B-120-treated PDO1, with corresponding log_2_(fold change) values. **c**, Gene Ontology enrichment analysis of differentially expressed genes in b. Dot size indicates gene number and color denotes −log_10_P value. Rich factor indicates the ratio of differentially expressed genes to the total number of genes in each term. **d**, PPI network linking DNA methylation, G1/S transition, DNA replication, one-carbon metabolism and senescence-associated cell-cycle arrest. **e**, **f**, Relative intracellular abundance of methionine (**e**) and homocysteine (**f**) in PDO1 following control or H3B-120 treatment, determined by targeted metabolomics. **g**, Hierarchically clustered heat map of methionine-cycle and related metabolites in PDO1 following H3B-120 treatment, with corresponding log_2_ fold changes shown on the right. **h**, Relative intracellular SAM abundance in Hepa1-6 cells, PDO1 and PDO2 following treatment with control, H3B-120, NH_4_Cl or H3B-120 plus 4-PBA. **i**, Quantification of relative SA-β-gal activity in PDO1 (left) and SA-β-gal-positive Hepa1-6 cells (right) following the indicated treatments. **j**, Representative SA-β-gal staining images of Hepa1-6 cells and PDO1 following treatment with control, H3B-120, NH_4_Cl or H3B-120 plus 4-PBA. Scale bars, 100 μm (j). Images are representative of three independent experiments. Data in e,f,h,i are mean ± s.d.; individual points represent biologically independent samples, with n = 3 per condition. Statistical significance was assessed using two-sided unpaired Student’s t-tests in e,f and one-way ANOVA followed by Tukey’s multiple-comparisons test in h,i. Exact P values are shown in the corresponding panels.

Immunofluorescence confirmed a marked reduction in MAT2A abundance in Hepa 1-6 cells and PDO1 after treatment with H3B-120 (Extended Data Fig. 1b). Targeted metabolomics in PDO1 further showed accumulation of methionine together with depletion of downstream homocysteine across the methionine cycle, one-carbon metabolism and related amino-acid pathways (Fig. 3e–g; Extended Data Fig. 1c). Consistent with restricted MAT2A-dependent methionine utilization, intracellular SAM was depleted in Hepa 1-6 cells, PDO1 and PDO2 following H3B-120 treatment (Fig. 3h). The combined accumulation of methionine and depletion of homocysteine and SAM therefore supported disruption of the MAT2A–SAM methyl-donor axis. To determine whether ammonia itself was sufficient to reproduce this metabolic state independently of pharmacological CPS1 inhibition, we exposed Hepa 1-6 cells, PDO1 and PDO2 to NH_4_Cl. Exogenous ammonia similarly reduced MAT2A abundance and depleted intracellular SAM, phenocopying the effects of CPS1 inhibition (Fig. 3h; Extended Data Fig. 1b). Conversely, lowering the ammonia burden with 4-PBA partially restored SAM abundance despite continued CPS1 inhibition (Fig. 3h), supporting ammonia accumulation as a direct contributor to disruption of MAT2A–SAM metabolism.

Because SAM provides the principal methyl donor for DNA methylation, we next examined whether its depletion was accompanied by changes in methylation-related and cell-cycle programs. H3B-120 reduced the expression of DNMT1, DNMT3A and DNMT3B, which mediate maintenance and de novo DNA methylation. This reduction in DNA methyltransferase expression, together with SAM depletion, indicated diminished cellular methylation capacity. In parallel, H3B-120 induced CDKN1A and CDKN2A, increased the expression of senescence-associated secretory phenotype genes and reduced LMNB1 expression (Fig. 3b–d). Immunofluorescence confirmed increased p21^CIP1^ and p16^INK4a^ abundance together with reduced LMNB1 abundance following H3B-120 treatment (Extended Data Fig. 1b, d). Senescence-associated β-gal (SA-β-gal) activity, a widely used marker of cellular senescence, was also markedly elevated in H3B-120-treated cells and PDOs (Fig. 3i, j). Direct exogenous ammonia recapitulated these molecular and phenotypic changes, whereas 4-PBA partially reduced SA-β-gal activity after H3B-120 treatment (Fig. 3i,j). Together, these findings place ammonia accumulation upstream of MAT2A–SAM disruption and senescence, and show that reducing the ammonia burden can partially reverse both the metabolic and phenotypic consequences of CPS1 blockade.

## Senescence reinforces ammoniagenesis to drive the ammonia–senescence metabolic ratchet towards ammonioptosis

Having established that ammonia retention promotes senescence, we next explored whether the senescent state reciprocally increases ammonia production. Senescent cells can acquire a GLS1-dependent glutaminolytic programme that supports their persistence^9, 18^. We therefore hypothesized that senescence-associated glutaminolysis could increase ammoniagenesis when ureagenic disposal is constrained, creating a self-reinforcing ammonia–senescence ratchet that progressively overwhelms intracellular ammonia handling (Fig. 4a). To examine this relationship independently of CPS1 inhibition, we induced senescence in Hepa1-6 cells and PDO1 with H_2_O_2_^19^. H_2_O_2_ markedly increased SA-β-gal staining in both models and was accompanied by increased GLS1 abundance and intracellular ammonia (Extended Data Fig. 2a–c). Thus, senescence induced independently of CPS1 blockade was associated with acquisition of a GLS1-high, ammonia-accumulating phenotype.

**Fig. 4.**
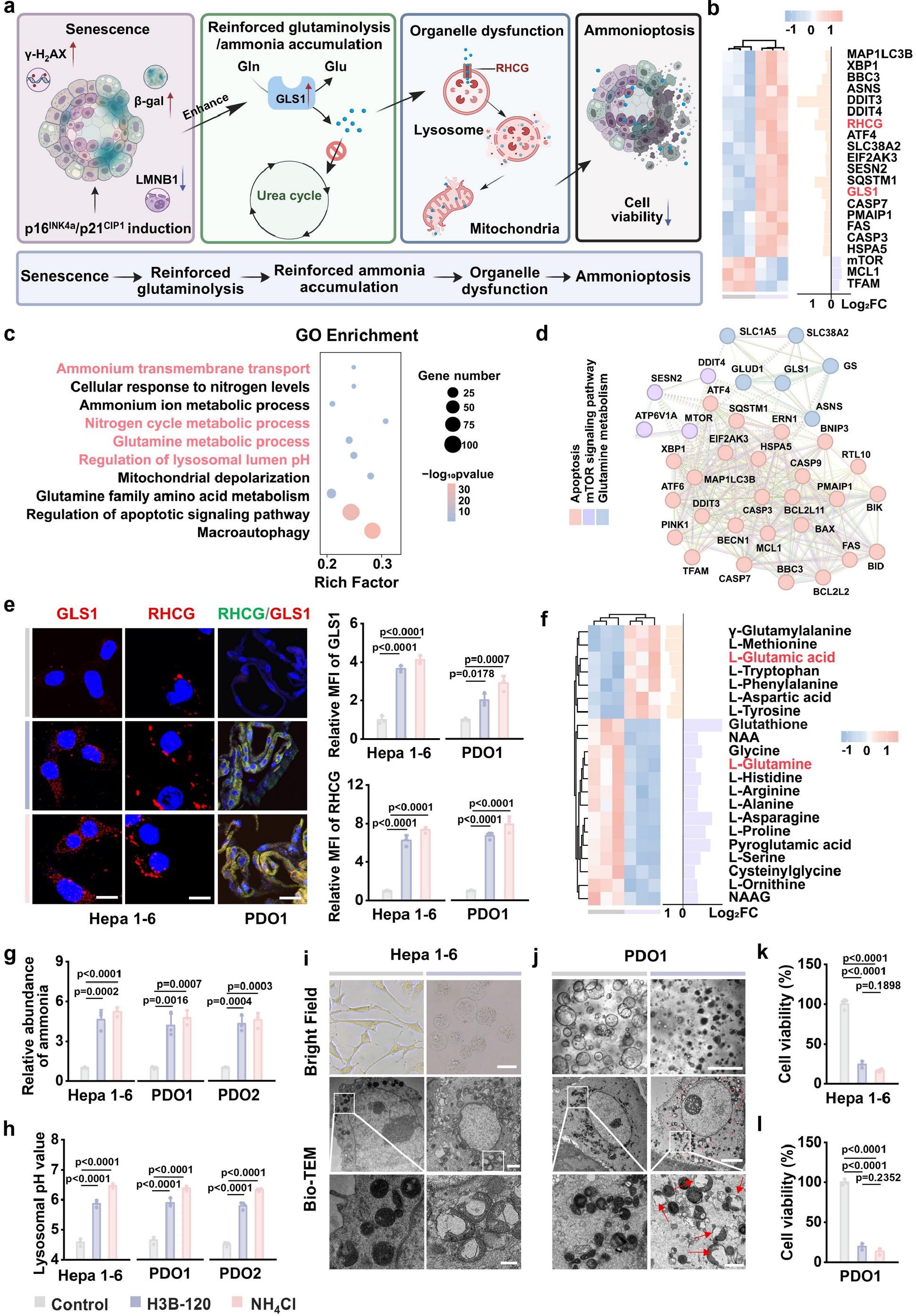
Senescence reinforces ammoniagenesis to drive the ammonia–senescence metabolic ratchet towards ammonioptosis. **a**, Schematic of the proposed ammonia–senescence ratchet linking senescence, enhanced glutaminolysis, ammonia accumulation, organelle dysfunction and ammonioptosis. **b**, Hierarchical clustering heat map of selected genes associated with glutamine metabolism, cellular stress and organelle homeostasis in PDO1 after H3B-120 treatment, with corresponding log_2_ fold-change values shown on the right. **c**, GO enrichment analysis of differentially expressed genes from the RNA-seq dataset in **b**. Dot size indicates gene number and color denotes −log_10_(P value). Rich factor indicates the ratio of differentially expressed genes to all genes annotated in each term. **d**, STRING protein–protein interaction network of differentially expressed genes associated with apoptosis, mTOR signalling and glutamine metabolism. Node colours denote the indicated functional categories. **e**, Representative GLS1 and RHCG immunofluorescence images and corresponding quantification of relative mean fluorescence intensity (MFI) in Hepa1-6 cells and PDO1 following treatment with control, H3B-120 or NH_4_Cl. **f**, Hierarchically clustered heat map of intracellular metabolites altered in PDO1 following H3B-120 treatment, with log_2_ fold changes shown on the right. **g**, **h**, Relative intracellular ammonia abundance (**g**) and lysosomal pH (**h**) in Hepa1-6 cells, PDO1 and PDO2 following treatment with control, H3B-120 or NH_4_Cl. **i**, **j**, Representative bright-field and transmission electron microscopy (TEM) images of Hepa1-6 cells (**i**) and PDO1 (**j**) following control or H3B-120 treatment. Red arrows indicate disrupted organelles. **k**, **l**, Cell viability of Hepa 1-6 cells (**k**) and PDO1 (**l**) after treatment with control, H3B-120 or NH_4_Cl. Scale bars, 20 μm (**e**); 20 μm, 2 μm and 500 nm from top to bottom in **i**; and 50 μm, 5 μm and 1 μm from top to bottom in **j**. Images are representative of three independent experiments. Data in **e,g,h,k,l** are mean ± s.d.; individual points represent biologically independent samples, with *n* = 3 per condition. Statistical significance was assessed using one-way ANOVA followed by Tukey’s multiple-comparisons test. Exact *P* values are shown in the corresponding panels.

We next examined whether CPS1 inhibition elicited the same metabolic phenotype. Transcriptomic analysis of H3B-120-treated PDO1 revealed coordinated changes in glutamine metabolism, ammonium transport and organelle-stress pathways (Fig. 4b). GO analysis identified enrichment of glutamine metabolism, nitrogen-cycle metabolism, ammonium transmembrane transport, lysosomal-pH regulation, mitochondrial depolarization, apoptotic signaling and macroautophagy (Fig. 4c; Extended Data Fig. 2d). PPI analysis grouped the differentially expressed genes into modules associated with glutamine metabolism, mTOR signaling, autophagy and apoptosis (Fig. 4d). Among the most prominent changes were increased expression of GLS1 and the ammonium transporter RHCG (Fig. 4b).

Immunofluorescence confirmed increased GLS1 and RHCG abundance in H3B-120-treated Hepa 1-6 cells and PDO1 (Fig. 4e). NH_4_Cl elicited a similar response, indicating that elevated ammonia itself can reinforce this metabolic programme. Targeted metabolomics further showed glutamine depletion together with glutamate accumulation following CPS1 inhibition, consistent with increased glutaminolytic activity (Fig. 4f; Supplementary Fig. 3). Importantly, intracellular ammonia measured after establishment of the senescence was higher than that observed during the earlier response to CPS1 inhibition across Hepa1-6 cells, PDO1 and PDO2 (Fig. 2i and Fig. 4g). This later increase coincided with GLS1 induction and the glutamine-to-glutamate shift. These findings support a biphasic model in which CPS1 inhibition initially increases ammonia by restricting ureagenic disposal, followed by a senescence-associated phase of reinforced glutaminolysis and further ammonia accumulation.

How this escalating ammonia burden affected organelle homeostasis was then examined. H3B-120 and NH_4_Cl increased lysosomal pH in Hepa 1-6 cells, PDO1 and PDO2 (Fig. 4h; Supplementary Fig. 4). In Hepa1-6 cells, lysosomal alkalinization was accompanied by reduced LysoTracker retention and increased lysosomal membrane permeabilization (Extended Data Fig. 2e,f). Mitochondrial membrane potential was also impaired, as indicated by loss of JC-1 aggregates and a reduced aggregate-to-monomer ratio (Extended Data Fig. 2g). Bright-field imaging showed cell rounding and loss of adherent morphology in Hepa1-6 cultures, whereas PDO1 exhibited shrinkage and loss of structural compactness (Fig. 4i,j). H&E staining further revealed disruption of organoid architecture and enlarged luminal or vacuole-like spaces (Extended Data Fig. 2h). Bio-TEM corroborated these changes, showing enlarged lysosome-like compartments, mitochondrial structural damage and extensive cytoplasmic vacuolization following CPS1 inhibition.

These organelle defects preceded loss of cell viability. Live/dead staining and Annexin V/PI flow cytometry showed increased mortality and expansion of dead-cell populations following CPS1 blockade or exogenous ammonia exposure (Extended Data Fig. 2i, j). Viability assays confirmed pronounced cell loss in Hepa 1-6 cells and multiple liver tumor PDOs (Fig. 4k, l; Extended Data Fig. 2k, l). Dose–response analysis further showed that H3B-120 caused a concentration-dependent reduction in Hepa 1-6 cell viability, whereas HUVEC viability was only modestly affected over the same concentration range (Extended Data Fig. 2m, n). These results indicate greater susceptibility of liver cancer cells than endothelial cells to CPS1 inhibition under the conditions tested. The secondary rise in ammonia that followed senescence was therefore accompanied by lysosomal buffering failure, mitochondrial injury and progression to ammonioptosis.

To determine whether this death response could be accounted for by canonical regulated cell-death pathways, we tested pharmacological inhibitors of apoptosis, ferroptosis and necroptosis. Z-VAD-fmk, ferrostatin-1 and necrostatin-1s did not significantly restore viability in H3B-120-treated Hepa1-6 cells or liver tumor PDOs (Extended Data Fig. 2o). In contrast, lowering the ammonia burden with 4-PBA produced a substantial partial rescue of viability across the same models (Extended Data Fig. 2o). Together with the accompanying lysosomal and mitochondrial injury, these results support an ammonia-dependent tumor-cell death programme in which senescence-associated metabolic reinforcement drives ammonia beyond the adaptive range and towards ammonioptosis.

## HBCG couples CPS1 blockade to metal-ion-mediated reinforcement of the ammonia–senescence ratchet

The preceding experiments identified restricted ureagenic disposal and senescence-associated ammoniagenesis as complementary nodes of the ammonia–senescence ratchet. We therefore engineered HBCG, an acid-responsive Co/Gd-doped layered double hydroxide nanoplatform carrying the CPS1 inhibitor H3B-120. Following cellular uptake, endolysosomal acidification was designed to release H3B-120 together with Co^2+^ and Gd^3+^. The LDH was designed to release H3B-120, Co^2+^ and Gd^3+^ after acid-triggered disassembly. H3B-120 restricts CPS1-dependent ammonia disposal, whereas Co^2+^ was incorporated to favor HIF-1α-associated glutaminolysis and Gd^3+^ to reinforce senescence-associated cell-cycle arrest. The basic LDH framework was further expected to weaken lysosomal acidification after endocytic uptake^20^. HBCG was thus designed to reinforce successive nodes of a single metabolic circuit coupling ammonia retention, senescence-associated ammoniagenesis and loss of organelle buffering^20^.

Structural and compositional analyses confirmed successful synthesis of CoGd-LDH. TEM revealed a uniform nanoscale morphology (Fig. 5a), while energy-dispersive X-ray elemental mapping showed the homogeneous spatial distribution of Al, Co, O and Gd throughout the nanoplatform, confirming uniform metal incorporation (Fig. 5b). X-ray diffraction (XRD) patterns matched the standard LDH reference profile, verifying formation of the characteristic layered double hydroxide structure (Fig. 5c). X-ray photoelectron spectroscopy (XPS) further resolved the expected elemental composition and chemical states of the doped metals (Fig. 5d-f). Inductively coupled plasma (ICP) analysis quantified Co and Gd contents at 20.3% and 4.6% respectively, providing independent compositional evidence for successful Co/Gd doping. Additionally, Rhodamine B-labeled CoGd-LDH showed time-dependent uptake and lysosomal accumulation in Hepa 1-6 cells (Extended Data Fig. 3a, b). Exposure to acidic conditions promoted LDH disassembly, as visualized by TEM, confirming lysosomal release of H3B-120 and incorporated metal ions following endocytic delivery (Extended Data Fig. 3c).

**Fig. 5.**
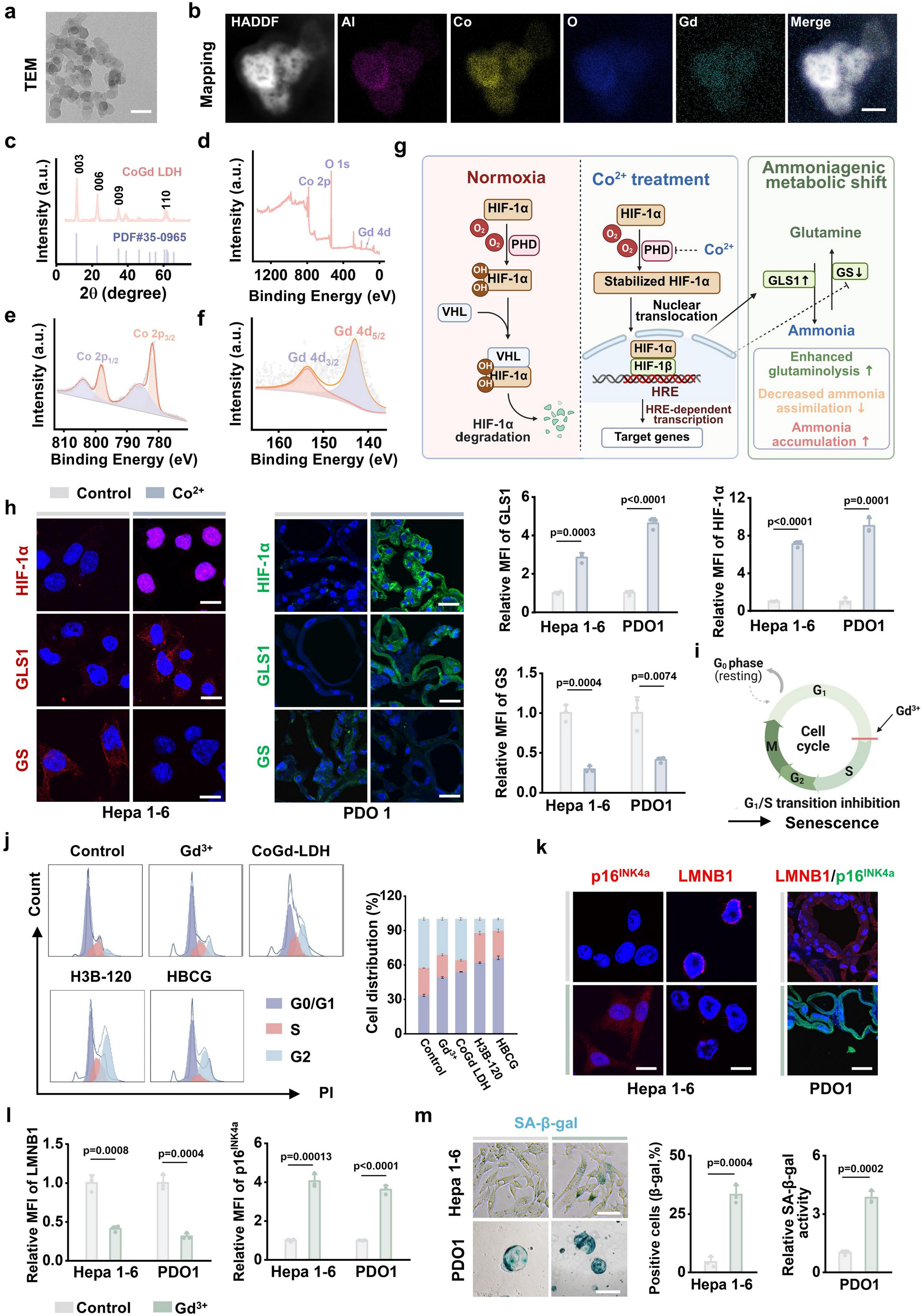
HBCG couples CPS1 blockade to metal-ion-mediated reinforcement of the ammonia– senescence ratchet. **a**, Representative transmission electron microscopy (TEM) image of CoGd-LDH. **b**, High-angle annular dark-field scanning transmission electron microscopy (HAADF-STEM) image and corresponding energy-dispersive X-ray spectroscopy elemental maps showing the spatial distribution of Al, Co, O and Gd in CoGd-LDH. **c**, X-ray diffraction pattern of CoGd-LDH showing the characteristic layered double hydroxide structure indexed to PDF#35-0965. **d–f**, X-ray photoelectron spectroscopy analysis of CoGd-LDH, showing the survey spectrum (**d**), Co 2p spectrum (**e**) and Gd 4d spectrum (**f**). **g**, Schematic of CoZ⁺-mediated stabilization of HIF-1α and the resulting shift towards enhanced glutaminolysis and reduced ammonia reassimilation. **h**, Representative immunofluorescence images and quantification of HIF-1α, GLS1 and GS in Hepa 1-6 cells and PDO1 treated with control or Co^2+^. Scale bars, 20 μm. **i**, Schematic of Gd3⁺-associated inhibition of G1/S cell-cycle progression and induction of senescence. **j**, Representative propidium iodide cell-cycle profiles and corresponding quantification of G0/G1-, S- and G2-phase distributions in Hepa1-6 cells following treatment with control, Gd^3+^, CoGd-LDH, H3B-120 or HBCG. **k**, **l**, Representative immunofluorescence images (**k**) and quantification (**l**) of LMNB1 and p16^INK4a^ in Hepa 1-6 cells and PDO1 treated with control or Gd^3+^. **m**, Representative SA-β-gal staining images and quantification of β-gal-positive Hepa 1-6 cells and relative SA-β-gal activity in PDO1 after control or Gd^3+^ treatment. Scale bars, 50 nm (a), 10 nm (b), 20 μm (h,k) and 100 μm (m). Images are representative of three independent experiments. Data in h,j, l,m are mean ± s.d.; individual points represent biologically independent samples, with n = 3 per condition. Statistical significance was assessed using two-sided unpaired Student’s t-tests for the two-group comparisons in h,l,m. Exact P values are shown in the corresponding panels.

The biological consequences of metal ions release were then examined directly. Co^2+^ can inhibit prolyl hydroxylase (PHD) activity (Fig. 5g), preventing hydroxylation-dependent recognition of HIF-1α by von Hippel–Lindau (VHL) E3 ubiquitin ligase complex and thereby stabilizing HIF-1α^21^. Co^2+^ treatment increased HIF-1α and GLS1 abundance while reducing glutamine synthetase (GS) abundance in Hepa 1-6 cells and PDO1 (Fig. 5h), which was consistent with increased glutamine catabolism and reduced ammonia reassimilation^22, 23^. Gd^3+^ was next examined for a potential senescence-reinforcing function in Hepa 1-6 cells and PDO1, by assessing canonical senescence-associated features, including G1/S cell-cycle arrest, SA-β-gal activity and senescence-marker expression. Flow-cytometric analysis showed that Gd^3+^ treatment induced pronounced accumulation of cells in the G0/G1 phase, accompanied by a corresponding reduction in the S phase population. HBCG further intensified this G1/S restriction, indicating that Gd^3+^ potentiates ammonia-induced senescence in the context of CPS1 blockade (Fig. 5i, j). This cell-cycle shift was accompanied by broader senescence phenotypes, as manifested by elevated SA-β-gal positivity and p16^INK4a^ abundance accompanied by LMNB1 loss. (Fig. 5k-m). HBCG therefore reinforces the ammonia– senescence ratchet at multiple nodes through enhanced ammoniagenesis, impaired detoxification, sustained senescence and disrupted lysosomal buffering, thereby progressively pushing senescent liver cancer cells toward ammonioptosis.

## HBCG drives a tumor-cell-biased ammonia–senescence ratchet towards ammonioptosis

The intact platform was then evaluated in Hepa 1-6 cells and liver tumor PDOs. Across Hepa 1-6 cells and PDOs, HBCG reduced urea production, increased HIF-1α protein abundance and suppressed GS abundance (Fig. 6a; Extended Data Fig. 4a). These findings indicated simultaneous impairment of CPS1-dependent urea-cycle flux and GS-mediated ammonia reassimilation, thereby redirecting cellular nitrogen handling toward intracellular ammonia retention. The resulting ammonia retention collapses methionine metabolism. HBCG strongly reduced MAT2A abundance in Hepa 1-6 cells and PDOs and depleted intracellular SAM, confirming that ammonia overload disrupts the MAT2A–SAM axis (Fig. 6b-d). Loss of methyl-donor homeostasis was accompanied by robust senescence induction in Hepa 1-6 cells and PDOs, as shown by elevated p16^INK4a^/p21^CIP1^ abundance, γ-H_2_AX and SA-β-gal activity together with repressed LMNB1 abundance (Extended Data Fig. 4b-d; Supplementary Fig. 5). Notably, HBCG induced a pronounced senescence-associated response (Fig. 6e), which exceeded those elicited by the clinically used liver cancer treatments tested^24–26^.

**Fig. 6.**
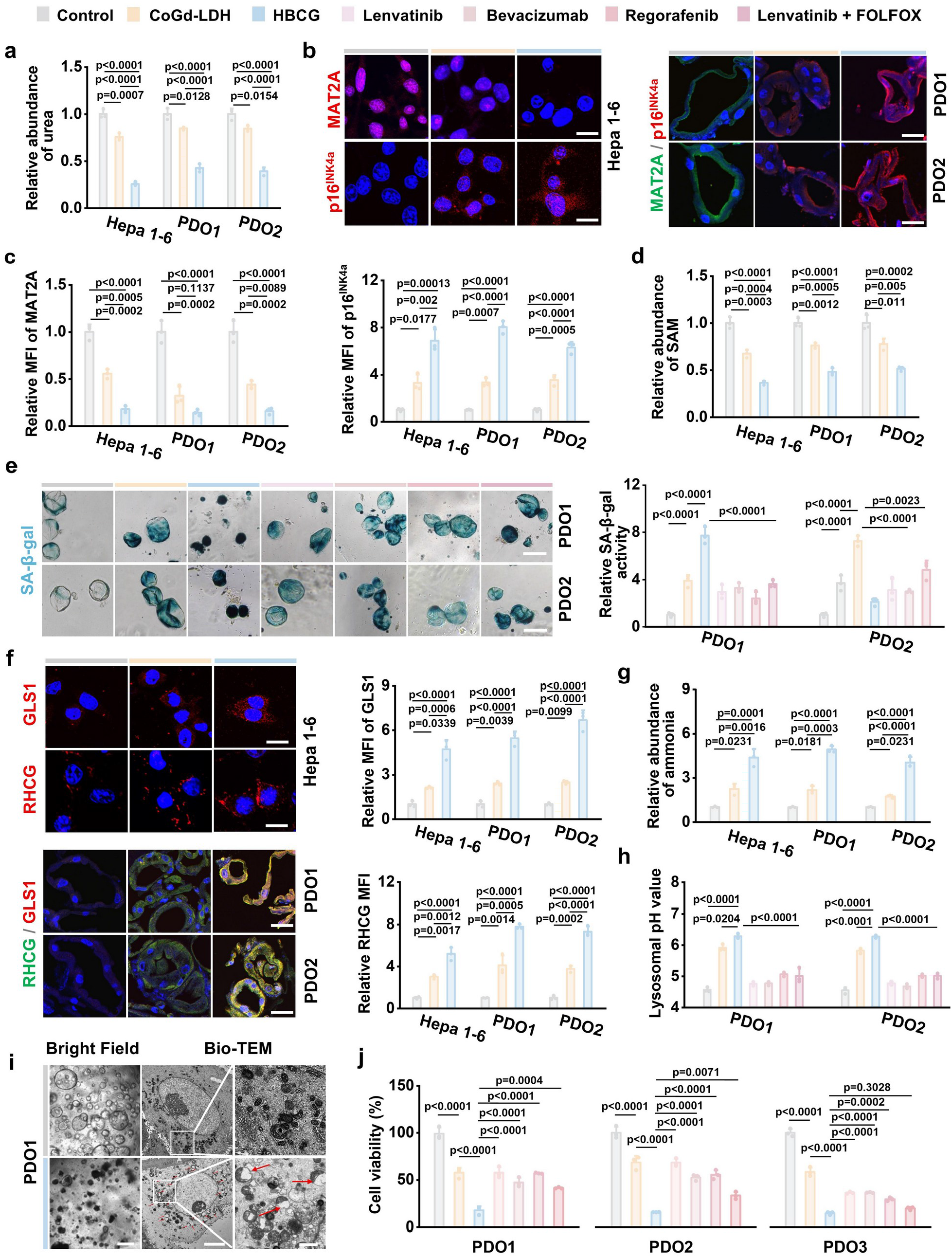
HBCG drives the ammonia–senescence ratchet towards ammonioptosis. **a**, Relative intracellular urea abundance in Hepa 1-6 cells, PDO1 and PDO2 after treatment with control, CoGd-LDH or HBCG**. b**, **c**, Representative MAT2A and p16^INK4a^ immunofluorescence images (**b**) and corresponding quantification of relative mean fluorescence intensity (MFI) (**c**) of in Hepa 1-6 cells, PDO1 and PDO2 after indicated treatment. **d**, Relative intracellular SAM abundance in Hepa 1-6 cells, PDO1 and PDO2 following the indicated treatments. **e**, Representative SA-β-gal staining images and corresponding quantification of relative SA-β-gal activity in PDO1 and PDO2 treated with control, CoGd-LDH, HBCG, lenvatinib, bevacizumab, regorafenib or lenvatinib plus FOLFOX. **f**, Representative GLS1 and RHCG immunofluorescence images and corresponding quantification of GLS1 and RHCG MFI in Hepa 1-6 cells, PDO1 and PDO2 after indicated treatment. **g**, Relative intracellular ammonia abundance in Hepa 1-6 cells, PDO1 and PDO2 following treatment with control, CoGd-LDH or HBCG. **h**, Lysosomal pH in PDO1 and PDO2 following treatment with control, CoGd-LDH, HBCG, lenvatinib, bevacizumab, regorafenib or lenvatinib plus FOLFOX. **i**, Representative bright field and Bio-TEM images of PDO1 following control or HBCG treatment. Red arrows indicate disrupted organelles. **j**, Cell viability of PDO1, PDO2 and PDO3 following treatment with control, CoGd-LDH, HBCG, lenvatinib, bevacizumab, regorafenib or lenvatinib plus FOLFOX. Nuclei in b and f were counterstained with DAPI. Scale bars, 20 μm (b,f), 100 μm (e), and 50 μm, 5 μm and 1 μm from left to right in i. Images are representative of three independent experiments. Data in a,c–h,j are mean ± s.d.; individual points represent biologically independent samples, with n = 3 per condition for each model. Statistical significance was assessed using one-way ANOVA followed by Tukey’s multiple-comparisons test in a,c–h,j. Exact P values are shown in the corresponding panels.

We further investigated whether senescent liver cancer cells adopt a GLS1-high ammoniagenic phenotype. Immunofluorescence showed elevated GLS1 abundance in HBCG-treated cells and PDOs, further reinforced by HIF-1α stabilization (Fig. 6f). This metabolic shift toward enhanced glutaminolysis, combined with blocked urea-cycle flux, imposed a substantial intracellular ammonia burden (Fig. 6g). Thus, HBCG converted senescence from an endpoint into a metabolically active state in which ammonia generation and detoxification failure became coupled. This distinguishes nonspecific cytotoxic stress from a feed-forward fate transition: the senescent program that limits proliferation also increases the nitrogen load that ultimately exceeds organellar buffering capacity, linking growth arrest to terminal death execution. This ammonia burden was further channeled towards lysosomal compartments through elevated RHCG-mediated transport. Escalating ammonia burden would favor proton consumption through NH_3_ protonation within acidic lysosomes, further potentiated by the intrinsic basicity of LDH, which progressively increased lysosomal pH (Fig. 6h; Extended Data Fig. 4e; Supplementary Fig. 6). This progressed to broader lysosomal dysfunction (Extended Data Fig. 4f, g). As lysosomal buffering became saturated, ammonia was retained in mitochondria, culminating in mitochondrial depolarization (Extended Data Fig.4h).

Bright-field and ultrastructural analyses corroborated this organelle dysfunction. HBCG-treated Hepa 1-6 cells exhibited cell rounding and loss of adherent morphology, whereas treated PDOs showed shrinkage, loss of epithelial compactness and architectural collapse. TEM further revealed cytoplasmic vacuolization, enlarged lysosome-like compartments and disruption of mitochondrial membranes and cristae (Fig. 6i; Extended Data Fig. 4i). Bright-field comparison with selected clinical agents and regimens provided an additional morphological benchmark: PDO1 organoids treated with bevacizumab, lenvatinib, regorafenib or unloaded CoGd-LDH largely retained discernible spheroid boundaries, whereas lenvatinib plus FOLFOX caused greater structural disruption; HBCG produced more pronounced organoid shrinkage and disintegration under the corresponding experimental conditions (Fig. 6i; Supplementary Fig. 7). These observations connect HBCG-induced ammonia accumulation with progressive lysosomal, mitochondrial and tissue-architectural damage.

The ensuing organellar damage caused by ammonia overwhelmed cellular homeostasis and culminated in cell death. Annexin V/PI flow cytometry showed a marked redistribution of HBCG-treated Hepa 1-6 cells from the viable population towards dead-cell fractions (Extended Data Fig. 4j, k), while live/dead staining independently confirmed an increase in PI-positive cells (Extended Data Fig. 4l). HBCG also reduced Hepa 1-6 viability more strongly than unloaded CoGd-LDH (Extended Data Fig. 4m). Benchmarking against lenvatinib, bevacizumab, regorafenib and FOLFOX plus lenvatinib, HBCG produced the strongest cytotoxic response across PDO1, PDO2 and PDO3 (Fig. 6j). Concentration–response analysis further showed a progressive loss of Hepa 1-6 viability with increasing HBCG concentrations, whereas HUVECs exhibited only a comparatively modest decline over the same concentration range (Extended Data Fig. 4n, Supplementary Fig. 8). Together with the preceding evidence of ammonia accumulation and lysosomal and mitochondrial injury, these findings support HBCG-induced ammonioptosis and indicate preferential susceptibility of liver tumor models to this metabolic intervention.

Ammonia exerts cell-type- and compartment-dependent effects on tumor immunity. Ammonia accumulation in the tumor microenvironment can impair effector T-cell function and weaken antitumor immunity. In contrast, Tregs can adapt to ammonia-rich conditions, shifting the immune balance towards a more suppressive state^11, 12, 27^. These studies suggest that indiscriminate ammonia elevation could damage effector T cells while favoring ammonia-adapted Tregs. Whether intracellular ammonia amplification by HBCG in tumor cells would translate into increased ammonia exposure of neighbouring immune cells was subsequently investigated. In tumor–PBMC co-cultures, HBCG markedly increased ammonia abundance in tumor cells, whereas ammonia in the shared culture medium increased only modestly and remained close to untreated group in CD8⁺ T cells (Extended Data Fig. 5a–c). By contrast, direct NH₄Cl treatment produced a broad extracellular ammonia load accompanied by increased CD8⁺ T-cell ammonia. Thus, HBCG preferentially confined the ammonia stress within tumor cells rather than distributing it broadly across the extracellular compartment. This restricted ammonia exposure was accompanied by preservation of T-cell viability and phenotype. CD8⁺ T-cell viability was largely maintained after HBCG treatment but substantially reduced by NH_4_Cl (Extended Data Fig. 5d). We next examined human T-cell responses to HBCG. HBCG largely preserved the frequencies of CD4⁺ T cells, Tregs, total CD8⁺ T cells and CD8⁺ effector T cells, with only a modest reduction in the GZMB⁺ CD8⁺ population. In contrast, NH_4_Cl caused broader perturbation of T-cell composition, reducing several effector populations while increasing the Treg fraction (Extended Data Fig. 5e–i). This difference is consistent with the distinct modes of ammonia exposure: HBCG relies on cellular uptake and intracellular metabolic amplification, whereas NH_4_Cl imposes an immediate and non-selective extracellular ammonia load.

## HBCG suppresses liver tumor growth in vivo by enforcing the ammonia– senescence ratchet

Having established that HBCG executes ammonioptosis in liver cancer cells and organoids, we explored whether it engaged the same fate trajectory in vivo. To test this, we established a syngeneic liver tumor model by subcutaneous implantation of Hepa1-6 cells into C57BL/6 mice and administered intratumorally on days 1, 3, and 5 after tumors reached 50–100 mm^3^, with tumor harvest on day 14 (Fig. 7a). Therapeutic responses diverged clearly across treatment groups. HBCG produced pronounced anti-tumor effects, with excised tumor volumes markedly smaller than untreated controls and comparable to NH_4_Cl-treated mice and sustained inhibition of tumor growth throughout treatment (Fig. 7b, c; Extended Data Fig. 6a, b). Tumor growth was consistently suppressed throughout treatment without overt systemic toxicity, as indicated by stable body weight across all groups (Fig. 7d).

**Fig. 7.**
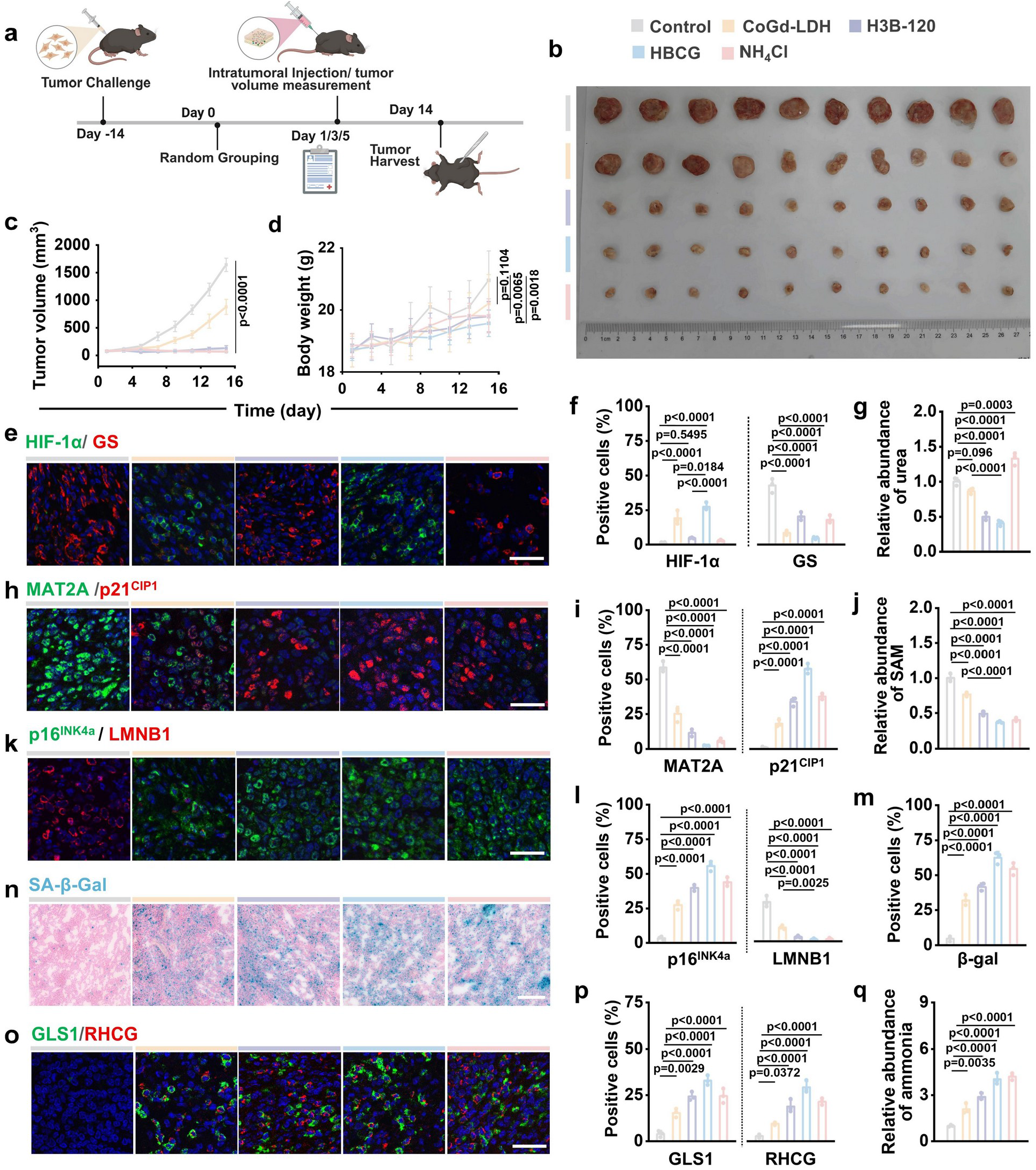
HBCG suppresses liver tumor growth in vivo by enforcing the ammonia–senescence ratchet. **a**, Schematic of the in vivo treatment schedule. C57BL/6 mice bearing subcutaneous Hepa1-6 tumors were randomized on day 0 and received intratumoral injections of the indicated treatments (control, CoGd-LDH, H3B-120, HBCG or NH_4_Cl) on days 1, 3 and 5; tumors were collected on day 14. **b**, Excised tumors from the indicated treatment groups on day 14. **c**, **d**, Tumor volume (c) and body weight (d) over time. **e**, **f**, Representative immunofluorescence images of HIF-1α (green) and GS (red) (**e**) and percentages of marker-positive cells (**f**). **g**, Relative intratumoral urea abundances after the indicated treatments. **h**, **i**, Representative immunofluorescence images of MAT2A (green) and p21^CIP1^ (red) (**h**) and percentages of marker-positive cells (**i**). **j**, Relative intratumoral SAM abundance. **k**, **l**, Representative immunofluorescence images of p16^INK4a^ (green) and LMNB1 (red) (k) and percentages of marker-positive cells (l). **m**, **n**, Percentage of SA-β-gal-positive cells (**m**) and representative SA-β-gal staining (**n**). **o**, **p**, Representative immunofluorescence images of GLS1 (green) and RHCG (red) (**o**) and percentages of marker-positive cells (**p**). **q**, Relative intratumoral ammonia abundance. Nuclei in e, h, k and o were counterstained with DAPI (blue). Scale bars, 50 μm (e, h, k, o) and 100 μm (n). Images are representative of three biologically independent tumors per group. For image quantification, each data point represents the mean value per tumor. Data are mean ± s.d.; n = 10 biologically independent mice per group (b–d) and n = 3 biologically independent tumors per group (f, g, i, j, l, m, p, q). c, d, two-way repeated-measures ANOVA with Tukey’s multiple-comparisons test; f, g, i, j, l, m, p, q, one-way ANOVA with Tukey’s multiple-comparisons test. Exact P values are shown.

To determine whether tumor control was accompanied by engagement of the ammonia– senescence ratchet, we analyzed tumor tissues across successive nodes of the proposed pathway. HBCG-treated tumors showed lower urea levels, increased HIF-1α abundance and reduced GS abundance (Fig. 7e–g). In the context of H3B-120-mediated CPS1 inhibition, these changes were consistent with restricted ureagenic ammonia disposal and diminished ammonia reassimilation (Fig. 7e-g). HBCG also reduced MAT2A abundance and intratumoral SAM levels (Fig. 7h–j). Suppression of the MAT2A–SAM methyl-donor axis was accompanied by increased p16^INK4a^ and p21^CIP1^ abundance, enhanced SA-β-gal activity and loss of LMNB1 (Fig. 7h, i,k–n), recapitulating the ammonia-associated senescence program observed in vitro.

The senescent state further hyperactivated GLS1-mediated glutaminolysis, reinforced by HIF-1α stabilization, thereby increasing ammonia production under conditions in which primary ureagenic disposal and compensatory GS-mediated reassimilation were both disabled (Fig. 7o-q). Concurrent upregulated RHCG redirected substantial ammonia toward lysosomal compartments, promoting organellar injury and driving tumor cells toward proliferative collapse and death, as evidenced by reduced Ki67 staining, robust TUNEL positivity and marked histopathological destruction in H&E staining (Extended Data Fig. 6c, d). These findings confirmed that escalating ammonia burden overwhelms lysosomal buffering, drives organellar injury, and culminates in ammonioptosis. Finally, we assessed systemic toxicity. H&E staining of major organs—heart, liver, spleen, lung, and kidney, showed no evident pathological injury, indicating that HBCG selectively targets tumor tissue without systemic toxicity (Extended Data Fig. 6e).

### Discussion

This work identifies intracellular ammonia as a cell-fate determinant that governs the establishment, maintenance and termination of tumor-cell senescence in liver cancer. Restricting CPS1-dependent ureagenic disposal drives intracellular ammonia accumulation, suppresses the MAT2A–SAM axis and promotes global DNA hypomethylation and senescence. The senescent state in turn enhances GLS1-dependent ammoniagenesis, establishing a self-reinforcing ratchet that progressively increases intracellular ammonia stress. Once ammonia exceeds lysosomal buffering capacity, lysosomal dysfunction and mitochondrial injury culminate in ammonioptosis, an ammonia-induced tumor-cell death. Thus, the senescence-sustaining metabolism can become self-destructive when driven beyond cellular tolerance, linking the establishment, maintenance and elimination of senescence within a single metabolic circuit.

This mechanism departs conceptually from conventional “one–two-punch” strategies, in which senescence creates state-specific vulnerabilities that are subsequently targeted by mechanistically distinct senolytic interventions^28^. Senescent cells, for example, rely on GLS1-dependent glutaminolysis for ammonia-mediated pH homeostasis, and GLS1 inhibition promotes senolysis by disrupting this support. Other senolytic approaches similarly target acquired survival dependencies, including apoptotic and ferroptotic vulnerabilities^8, 9, 18^. Co-delivery can improve spatial and pharmacokinetic coordination, but does not remove this mechanistic separation between senescence induction and clearance^1, 29, 30^. Here, we reverse this logic: maintaining GLS1-dependent glutaminolysis while restricting ammonia disposal turns a senescence-supporting metabolic output into a toxic intracellular burden. The same metabolic program that sustains senescence therefore drives its elimination, without a separate senolytic intervention. Senescence therefore functions not simply as an endpoint or a vulnerability-generating state, but as an active metabolic amplifier toward death.

Our results further refine the emerging view of ammonia as a context-dependent metabolite in cancer. Tumor-associated ammonia can be reassimilated to support anabolism, whereas its accumulation in the tumor microenvironment can impair antitumor T-cell function and favor an immunosuppressive state^31–34^. Excessive intracellular ammonia can also induce lysosomal alkalinization, mitochondrial injury and death in effector CD8^+^ T cells^12^. Thus, ammonia is neither intrinsically tumor-promoting nor tumor-toxic; its effects depend on cellular exposure and the capacity for reassimilation, disposal and organelle buffering. The therapeutic objective is therefore not generalized ammonia elevation, but selective overloading of ammonia handling within tumor cells. HBCG exploits this distinction by restricting CPS1-dependent ammonia disposal and reinforcing senescence-associated ammoniagenesis without introducing an exogenous ammonia-generating reaction^35–37^. By coupling CPS1 inhibition with reinforcement of senescence-associated ammoniagenesis and impaired organelle buffering, HBCG preferentially increased ammonia stress in tumor cells. In the tumor cell–PBMC co-culture system, this was accompanied by only a modest rise in extracellular ammonia and little increase in intracellular ammonia in CD8⁺ T cells, whereas direct NH_4_Cl exposure increased ammonia more broadly. These findings support a tumor-cell-biased rather than indiscriminate amplification of ammonia stress, although they do not establish absolute tumor specificity or systemic immune safety. Extracellular ammonia depletion and tumor-cell-intrinsic ammonia amplification are therefore not opposing strategies: the former limits microenvironmental ammonia exposure, whereas the latter seeks to exceed ammonia-handling capacity selectively within tumor cells. Their therapeutic compatibility will ultimately depend on achieving sufficient compartmental selectivity.

Several boundaries remain to be defined. First, the generality of the ammonia–senescence ratchet will likely depend on tumor-specific differences in glutamine utilization, ammonia production and disposal, reassimilation, and organelle-buffering capacity. Second, the current population-level measurements do not establish that individual cells continuously move from adaptation through senescence to ammonioptosis; longitudinal lineage tracing combined with single-cell metabolic and epigenetic profiling will be needed to distinguish a true directional trajectory from divergent cell-fate branches. Finally, the therapeutic window remains to be defined in normal hepatocytes and in immune populations in vivo, particularly under systemic delivery conditions.

Together, our findings show that a metabolic program sustaining senescent-cell survival can drive its elimination when pushed beyond cellular tolerance. The ammonia–senescence ratchet thereby links senescence induction, maintenance and cell-autonomous death within the same metabolic process, while providing a framework for selectively exploiting tumor-cell-intrinsic ammonia stress.

## Methods

### Small molecules and chemicals

H3B-120 (SJ-MX0225) was purchased from SparkJade. CoCl_2_·6H_2_O, GdCl_3_·6H_2_O and NH_4_Cl were obtained from Macklin. Lenvatinib (HY-10981), bevacizumab (HY-P9906A), regorafenib (HY-10331), 5-fluorouracil (HY-90006) and oxaliplatin (HY-17371) were obtained from MedChemExpress (Monmouth Junction, NJ, USA).

### Cell-culture reagents and consumables

Dulbecco’s modified Eagle’s medium (DMEM; JYK-SJ-003) was purchased from Jin Yuan Kang Biotechnology. RPMI-1640 medium (C11875500BT) was obtained from Gibco. Penicillin– streptomycin (KGL2303) was obtained from Keygen BioTECH. Matrigel (abs9495) was obtained from Absin. Phosphate-buffered saline (PBS) was purchased from Wuhan Procell Biotechnology Co., Ltd. (Wuhan, China). Serum-free cell-freezing medium (S9050S) was obtained from Uelandy (Suzhou, China), and Mycoplasma Elimination Reagent (C30200) was obtained from New Cell & Molecular Biotech. Human Peripheral Blood Lymphocyte Isolation Solution (Ficoll-based; P8900) was purchased from Solarbio. RNAkey Total RNA Extraction Reagent (chloroform substitute; SM139-02) was purchased from Seven/Abcells (Beijing, China). Recombinant human IL-2 (200-02-100UG; PeproTech, Thermo Fisher Scientific) was used for T-cell culture. Cell-culture dishes and plates, round coverslips and 20-mm glass-bottom dishes were obtained from NEST Biotechnology Co., Ltd. (Wuxi, China). Ultrafiltration centrifugal devices were obtained from Cobetter, and 96-well microplates were obtained from Bioland.

### Assay kits, stains and fluorescent probes

The CytoDet JC-1 Mitochondrial Membrane Potential Assay Kit was obtained from Accurate Biotechnology (Hunan) Co., Ltd. (Changsha, China). The Live and Dead Cell Double Staining Kit was purchased from Abbkine (Wuhan, China). The Annexin V-FITC/PI Apoptosis Detection Kit (RK05875) was obtained from ABclonal. The PI-based Cell Cycle and Apoptosis Analysis Kit (HY-K1071; MedChemExpress, Monmouth Junction, NJ, USA) was used for cell-cycle analysis. Thiazolyl Blue (KM9350) was obtained from KKL MED. DAPI (abs47047616) was obtained from Absin. Lyso-Tracker Green (IF1850) and the AO Fluorescent Stain Kit (CA1142) were purchased from Solarbio. LysoTracker Red and LysoSensor Yellow/Blue DND-160 (PDMPO) were obtained from Yeasen. LysoSensor Yellow/Blue DND-160 was used to measure lysosomal pH. The SA-β-gal Staining Kit was obtained from Servicebio. The DNA Damage Detection Kit for γ-H2AX immunofluorescence was purchased from Beyotime Biotechnology. The Ammonia Assay Kit (ab83360) was purchased from Abcam. The Urea Content Assay Kit was purchased from Beijing Boxbio Science & Technology Co., Ltd. The SAM ELISA kit was obtained from AMOU LUNCHANGSHUO BIOTEC, CO., LTD. All staining and biochemical assay kits were used according to the manufacturers’ instructions. Cyto-Fast^TM^ Fix/Perm Buffer Set (426803), True-Nuclear^TM^ Transcription Factor Buffer Set (424401), Human TruStain FcX^TM^ (422301) and Cell Staining Buffer (420201) were obtained from BioLegend.

### Antibodies

The following primary antibodies were used: mouse monoclonal anti-LMNB1 antibody (ELK Biotechnology); anti-p21^CIP1^ antibody (R381102; ZenBio); rabbit anti-CPS1 monoclonal antibody (A68975; Nature Biosciences); rabbit monoclonal anti-GLS1 antibody (A23189; ABclonal); anti-p16^INK4a^ antibody (ab211542; Abcam); rabbit polyclonal anti-GS antibody (100715-T44; Sino Biological Inc.); rabbit polyclonal anti-MAT2A antibody (55309-1-AP; Proteintech); rabbit polyclonal anti-RHCG antibody (bs-7638R; Bioss); and HIF-1α Antibody (HY-P84619; MedChemExpress, Monmouth Junction, NJ, USA). Anti-CD3 (E-AB-F10010) and anti-CD28 (E-AB-F11950) activating antibodies were obtained from Elabscience Biotechnology Co.,Ltd. The following fluorophore-conjugated antibodies were used for flow cytometry: PerCP anti-human CD45 (368505; BioLegend); FITC anti-human CD3 (300305; BioLegend); APC/Cyanine7 anti-human CD4 (317417; BioLegend); PE anti-human FOXP3 (320207; BioLegend); Spark Red™ 718 anti-human CD25 (356157; BioLegend); APC/Cyanine7 anti-human CD8a (301015; BioLegend); PE/Cyanine7 anti-human/mouse Granzyme B Recombinant (372213; BioLegend); PE anti-human CD197 (CCR7) (353203; BioLegend); and APC anti-human CD45RA (304111; BioLegend)

### Material synthesis and characterization

CoAl-LDH nanoparticles were first prepared by alkaline co-precipitation of Co^2+^ and Al^3+^ precursors at a Co:Al molar ratio of 2:1, followed by hydrothermal treatment. The resulting CoAl-LDH nanoparticles were then isolated, redispersed in an aqueous Gd^3+^ solution and subjected to a second hydrothermal treatment to introduce Gd^3+^ into the preformed LDH nanoparticles, yielding Gd-modified CoAl-LDH nanoparticles, hereafter referred to as CoGd-LDH. H3B-120 was subsequently loaded onto the CoGd-LDH nanoparticles to obtain HBCG.

The pH-dependent release of H3B-120 from HBCG was evaluated in media at pH 7.4 or 5.5. Released H3B-120 was quantified by ultraviolet–visible spectroscopy. Particle morphology, elemental distribution, crystal structure and surface chemical states were characterized by TEM, Mapping, XRD and XPS, respectively.

### Cell culture

Hepa 1-6 mouse hepatocellular carcinoma cells and human umbilical vein endothelial cells (HUVECs) were obtained from the Shanghai Institute of Biochemistry and Cell Biology, Chinese Academy of Sciences. Cells were maintained in high-glucose DMEM supplemented with 10% fetal bovine serum and 1% penicillin–streptomycin. Cultures were maintained at 37 °C in a humidified atmosphere containing 5% CO_2_.

### Patient-derived liver cancer organoids

Patient-derived organoids were established from surgically resected liver tumor specimens after obtaining written informed consent. PDO1 and PDO3 were derived from hepatocellular carcinoma, whereas PDO2 was derived from intrahepatic cholangiocarcinoma. Tumor tissues were mechanically dissociated, enzymatically digested and filtered before red blood cell lysis. The resulting cells were embedded in Matrigel and maintained in liver cancer organoid culture medium. Organoids were cultured at 37 °C under 5% CO_2_, with medium replacement every three days. The study was approved by the Ethics Committee of Shandong Provincial Hospital Affiliated to Shandong First Medical University.

### Isolation and ex vivo activation of peripheral blood mononuclear cells (PBMCs) from liver tumor patients

Peripheral blood mononuclear cells (PBMCs) were isolated from peripheral blood samples obtained from patients with hepatocellular carcinoma by Ficoll density-gradient centrifugation at 2,000 r.p.m. for 20 min at room temperature. Cells collected from the plasma–Ficoll interface were washed three times with PBS and resuspended at 1.0 × 10^6^ cells/mL in RPMI-1640 medium supplemented with anti-CD3 antibody (3 μg/mL), anti-CD28 antibody (1.5 μg/mL), recombinant human IL-2 (500 U/mL), 10% fetal bovine serum and 1% penicillin–streptomycin. PBMCs were cultured for four days at 37 °C in a humidified atmosphere containing 5% CO_2_ before subsequent treatment.

### Cellular uptake and lysosomal localization

Cellular uptake was evaluated by incubating Hepa 1-6 cells with RhB-labelled CoGd-LDH for the indicated durations. Cells were subsequently washed, fixed and examined by confocal laser-scanning microscopy. For lysosomal co-localization analysis, cells were stained with LysoTracker after exposure to RhB-labelled CoGd-LDH. Cellular fluorescence intensity and lysosomal co-localization were quantified using ImageJ.

### In vitro treatments and cell-viability assays

Hepa 1-6 cells, HUVECs and patient-derived organoids were treated with NH_4_Cl, H3B-120, CoGd-LDH, HBCG or the indicated clinical comparator drugs. Cell viability was measured by MTT assay and normalized to the corresponding untreated control. Cell viability and death were further assessed by Calcein-AM/PI staining. Apoptosis was quantified by Annexin V-FITC/PI flow cytometry. Cell-cycle distributions were determined by PI/RNase staining followed by flow-cytometric analysis.

### Measurement of ammonia, urea and SAM

Intracellular ammonia, urea and S-adenosylmethionine levels were quantified using commercial assay kits according to the manufacturers’ instructions. Hepa 1-6 cells or PDOs were treated with the indicated formulations, collected after treatment and processed for biochemical measurements. Values were normalized to cell number, protein content or sample amount, as appropriate.

Activated PBMCs were co-cultured with tumor cells and treated with the indicated formulations. After treatment, tumor cells and CD8⁺ T cells were separated by flow-cytometric sorting and collected for intracellular ammonia measurement, while culture supernatants were collected in parallel for extracellular ammonia analysis.

### Senescence, lysosomal and mitochondrial assays

Senescence was evaluated by SA-β-gal staining and immunofluorescence analysis of p16^INK4a,^ p21^CIP1^ and LMNB1. Lysosomal pH was measured using LysoSensor Yellow/Blue DND-160, and relative lysosomal alkalinization was calculated from the fluorescence intensity ratio. Lysosomal membrane permeabilization was assessed by acridine orange staining, with loss of red punctate lysosomal fluorescence and increased diffuse green fluorescence used as indicators of lysosomal leakage. Mitochondrial membrane potential was determined using JC-1 staining and quantified as the red-to-green fluorescence intensity ratio. Fluorescence images were acquired by fluorescence microscopy or confocal microscopy and quantified using ImageJ.

### Immunofluorescence staining

Cells and organoids were fixed with paraformaldehyde, permeabilized and blocked before incubation with the indicated primary antibodies. Samples were subsequently incubated with fluorophore-conjugated secondary antibodies, and nuclei were counterstained with DAPI. Double immunofluorescence staining was performed on organoids and tumor sections where indicated. Images were acquired by confocal laser-scanning microscopy, and mean fluorescence intensity was quantified using ImageJ.

### Transcriptomic and targeted metabolomic analyses

PDO1 organoids were treated with H3B-120, with untreated organoids serving as controls. RNA sequencing and targeted liquid chromatography–tandem mass spectrometry were performed by Novogene Co., Ltd. Differentially expressed genes were subjected to Gene Ontology and Kyoto Encyclopedia of Genes and Genomes enrichment analyses. Targeted metabolomics quantified metabolites associated with ammonia metabolism, the urea cycle, glutamine–glutamate metabolism and the methionine cycle. Detailed experimental and analytical procedures are provided in the Supplementary Methods.

### In vivo anti-tumor study

Male C57BL/6 mice were used to establish subcutaneous Hepa 1-6 tumors. Hepa 1-6 cells were implanted subcutaneously into the right flank of each mouse. When tumor volumes reached approximately 50–100 mm^3^, mice were randomly assigned to five groups: control, NH_4_Cl, H3B-120, CoGd-LDH and HBCG. The indicated formulations were administered by intratumoral injection according to the experimental schedule. Tumor volumes and body weights were monitored throughout treatment. At the experimental endpoint, tumors and major organs were collected for histological, immunofluorescence and biochemical analyses. Animal experiments complied with the Chinese National Regulations for the Management and Use of Laboratory Animals. All procedures were approved by the relevant institutional animal ethics committee.

### Statistical analysis

Quantitative data are presented as the mean ± s.d. Individual data points represent biological replicates or individual animals, as defined in the figure legends. Statistical analyses were performed using GraphPad Prism.

Two-group comparisons were performed using two-sided unpaired Student’s t-tests. Multiple groups were compared by one-way ANOVA followed by Tukey’s multiple-comparisons test. Tumor-growth and body-weight trajectories were analyzed by two-way ANOVA followed by the specified multiple-comparisons test.

Exact P values and sample sizes are reported in the corresponding figures and legends. A two-sided P < 0.05 was considered statistically significant.

#### Data availability

All data supporting the findings of this study have been included in the article and its Supplementary Information. Source data are provided with this paper.

## Acknowledgements

This project was funded by Advanced Materials-National Science and Technology Major Project (2025ZD0617100), the National Natural Science Foundation of China (52272156), the Qingdao Science and Technology Benefiting the People Demonstration Project (25-1-5-smjk-15-nsh) and Taishan Scholar Program of Shandong Province. All PDO experiments were approved by the Ethics Committee of Shandong Provincial Hospital Affiliated to Shandong First Medical University (SWYX: NO.2026-517), and written informed consent was obtained from all patients before sample collection. All animal experiments were approved by the Ethical Committee of Shandong University under production license number SCXK 2019-0001 and use license number SYXK 2019-0005. The authors would like to thank Xiaoju Li, Haiyan Sui and Xueyun Geng for their help with the TEM as well as Sen Wang, Haiyan Yu, Xiaomin Zhao and Yuyu Guo for their kind help with flow cytometry analysis from Core Facilities for Life and Environmental Sciences at the SKLMT (State Key Laboratory of Microbial Technology), Shandong University.

## Author contributions

Y.H. designed the study, synthesized the materials, performed most in vitro and in vivo experiments, analyzed the data and drafted the manuscript. M.H. performed validation experiments in patient-derived organoids and contributed to data analysis. J.W. provided three independent PDO models. Y.X. provided experimental guidance. S.Z. assisted with cell-cycle and flow-cytometry experiments. C.L., L.H. and Y.Z. supervised the project, secured funding, guided the experimental design and revised the manuscript. All authors discussed the results and approved the final manuscript.

---

**Extended Data Fig. 1.**
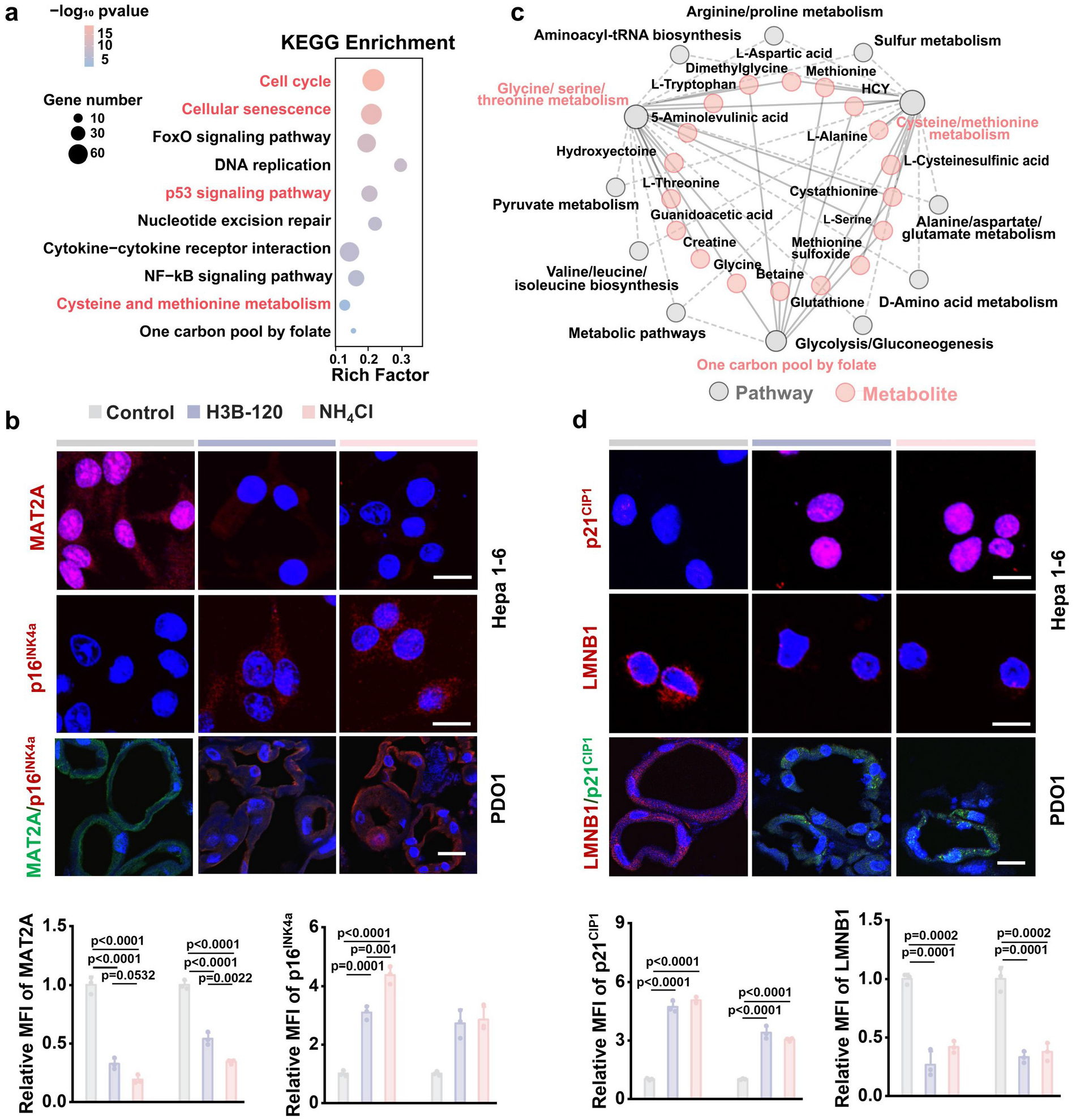
Ammonia stress suppresses MAT2A-dependent methyl-donor metabolism and induces senescence in liver tumor models. **a**, Kyoto Encyclopedia of Genes and Genomes (KEGG) pathway enrichment analysis of differentially expressed genes identified from the RNA-seq dataset shown in Fig. 3b following H3B-120 treatment in PDO1. Dot size indicates gene number, colour denotes −log_10_(P value), and rich factor represents the proportion of differentially expressed genes assigned to each pathway. **b**, Representative immunofluorescence images and corresponding quantification of MAT2A and p16^INK4a^ in Hepa1-6 cells and PDO1 following treatment with control, H3B-120 or NH_4_Cl. **c**, Integrated metabolite–pathway network linking differentially abundant metabolites to enriched metabolic pathways following H3B-120 treatment in PDO1. Pink nodes indicate metabolites and grey nodes indicate metabolic pathways. **d**, Representative immunofluorescence images and corresponding quantification of p21^CIP1^ and LMNB1 in Hepa1-6 cells and PDO1 following treatment with control, H3B-120 or NH_4_Cl. Nuclei in b and d were counterstained with DAPI. Scale bars, 20 μm (b,d). Images are representative of three independent experiments. Quantitative data in b,d are mean ± s.d.; individual points represent biologically independent samples, with n = 3 per condition. Statistical significance was assessed using one-way ANOVA followed by Tukey’s multiple-comparisons test in b,d. Exact P values are shown in the corresponding panels.

**Extended Data Fig. 2.**
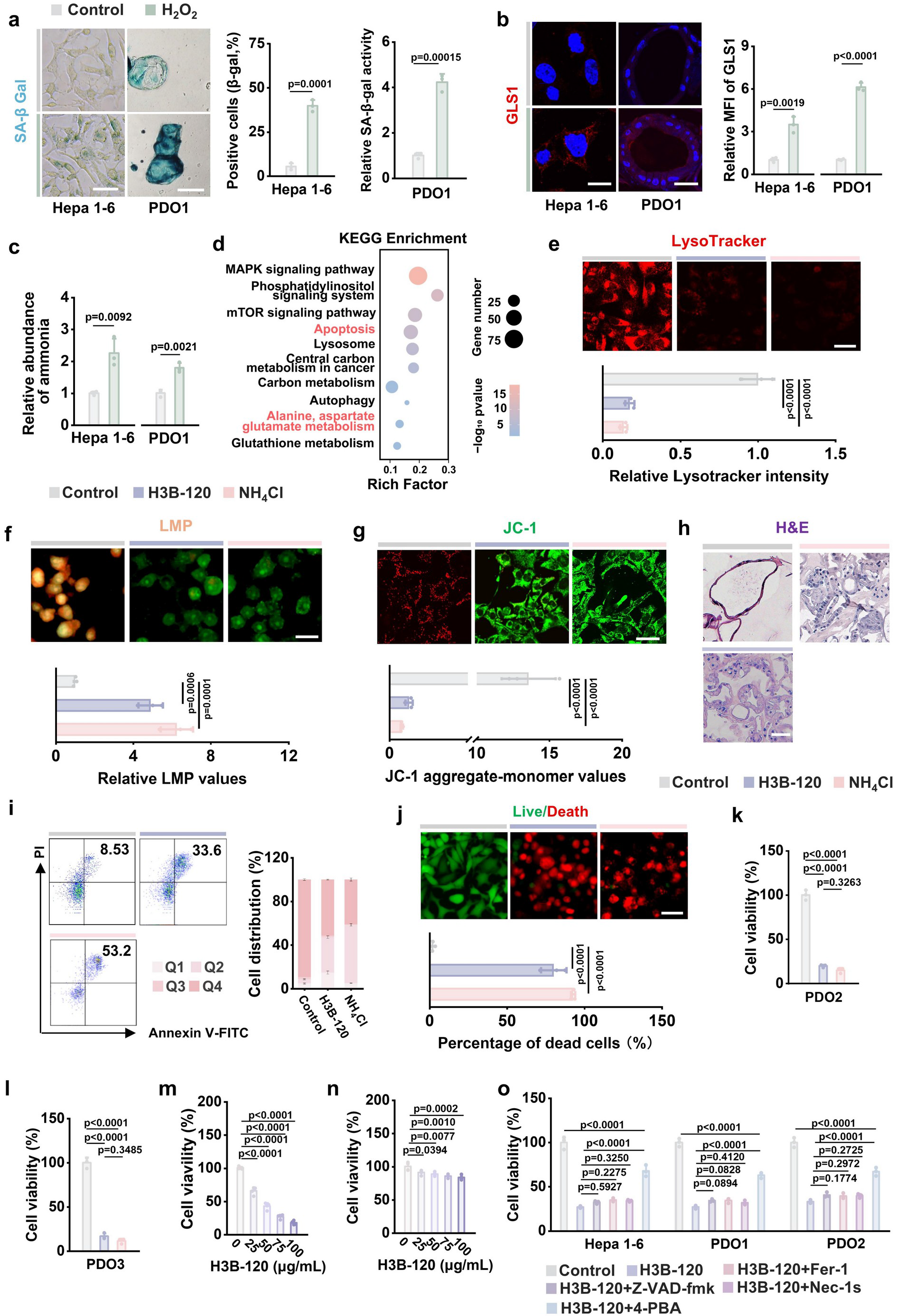
Senescence-associated ammoniagenesis accompanies organelle dysfunction and cell death in liver tumor models. **a**, Representative SA-β-gal staining and quantification of SA-β-gal-positive Hepa1-6 cells and relative SA-β-gal activity in PDO1 after control or H_2_O_2_ treatment. **b**, Representative GLS1 immunofluorescence images and corresponding quantification of relative mean fluorescence intensity (MFI) in Hepa1-6 cells and PDO1 following control or H_2_O_2_ treatment. Nuclei were counterstained with DAPI. **c**, Relative intracellular ammonia abundance after the indicated treatments. **d**, Kyoto Encyclopedia of Genes and Genomes (KEGG) pathway enrichment analysis of differentially expressed genes in H3B-120-treated PDO1 relative to untreated controls. Dot size indicates gene number, colour denotes −log_10_(P value), and rich factor represents the proportion of differentially expressed genes assigned to each pathway. **e**, Representative LysoTracker images and quantification of fluorescence intensity in Hepa1-6 cells treated with control, H3B-120 or NH_4_Cl. **f**, Representative fluorescence images and quantification of lysosomal membrane permeabilization (LMP) in Hepa 1-6 cells after the indicated treatments. **g**, Representative JC-1 fluorescence images and quantification of the aggregate-to-monomer fluorescence ratio in Hepa 1-6 cells after the indicated treatments. **h**, Representative H&E-stained sections of liver tumor PDOs after the indicated treatments. **i**, Representative Annexin V–FITC/PI flow-cytometry plots and corresponding quantification of cell distributions among the four quadrants in Hepa1-6 cells following the indicated treatments. **j**, Representative live/dead staining images and corresponding quantification of dead Hepa 1-6 cells after the indicated treatments. **k**, **l**, Viability of PDO2 (**k**) and PDO3 (**l**) after treatment with control, H3B-120 or NH_4_Cl. **m**, **n**, Concentration-dependent effects of H3B-120 on the viability of Hepa1-6 cells (m) and human umbilical vein endothelial cells (HUVECs; n). **o**, Cell viability of Hepa1-6 cells, PDO1 and PDO2 following treatment with control, H3B-120, H3B-120 plus Z-VAD-fmk, H3B-120 plus ferrostatin-1 (Fer-1), H3B-120 plus necrostatin-1s (Nec-1s), or H3B-120 plus 4-PBA. Scale bars, 100 μm (a), 20 μm (b,e–g,j) and 50 μm (h). Images are representative of three independent experiments. Data in a– c,e–g,j–o are mean ± s.d.; individual points represent biologically independent samples, with n = 3 per condition. Statistical significance was assessed using two-sided unpaired Student’s t-tests in a–c and one-way ANOVA followed by Tukey’s multiple-comparisons test in e–g,j–o. Exact P values are shown in the corresponding panels.

**Extended Data Fig. 3.**
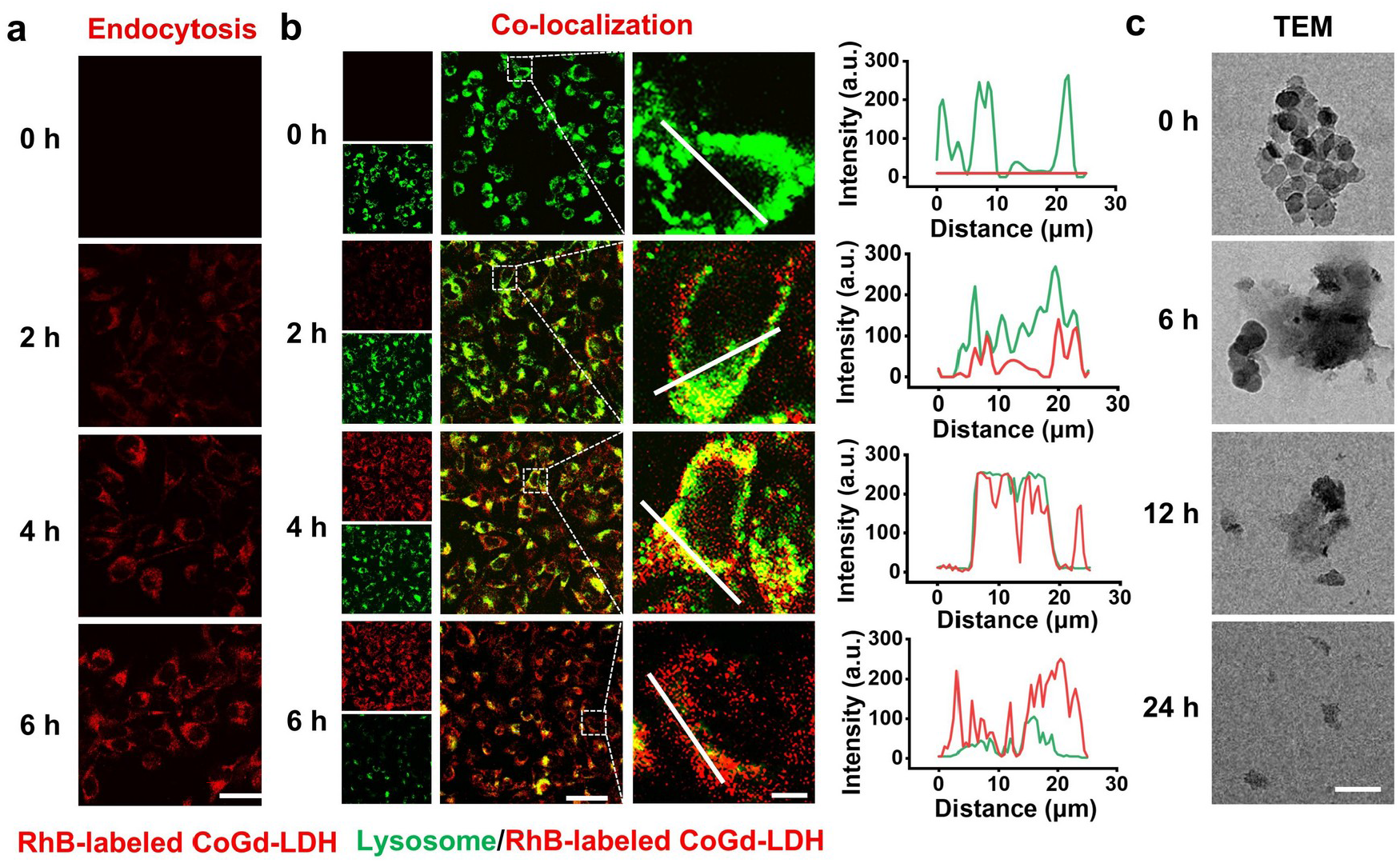
Time-dependent cellular uptake, lysosomal trafficking and dissociation of CoGd-LDH. **a**, **b**, Time-dependent confocal fluorescence imaging of cellular endocytosis (**a**) and lysosomal co-localization (**b**) of RhB-labeled CoGd-LDH nanoparticles in Hepa 1-6 cells over 6 h. CoGd-LDH is shown in red and lysosomes in green. Co-localization analysis in **b** shows merged images, magnified views of the boxed regions and corresponding line-scan fluorescence intensity profiles along the indicated white lines. Scale bars, 20 μm (**a**), 50 μm (**b**, main images) and 5 μm (**b,** magnified images). **c**, Representative TEM images showing time-dependent morphological dissociation of CoGd-LDH nanoparticles after incubation for 0, 6, 12 and 24 h. Scale bar, 100 nm. Representative images from three independent experiments are shown.

**Extended Data Fig. 4.**
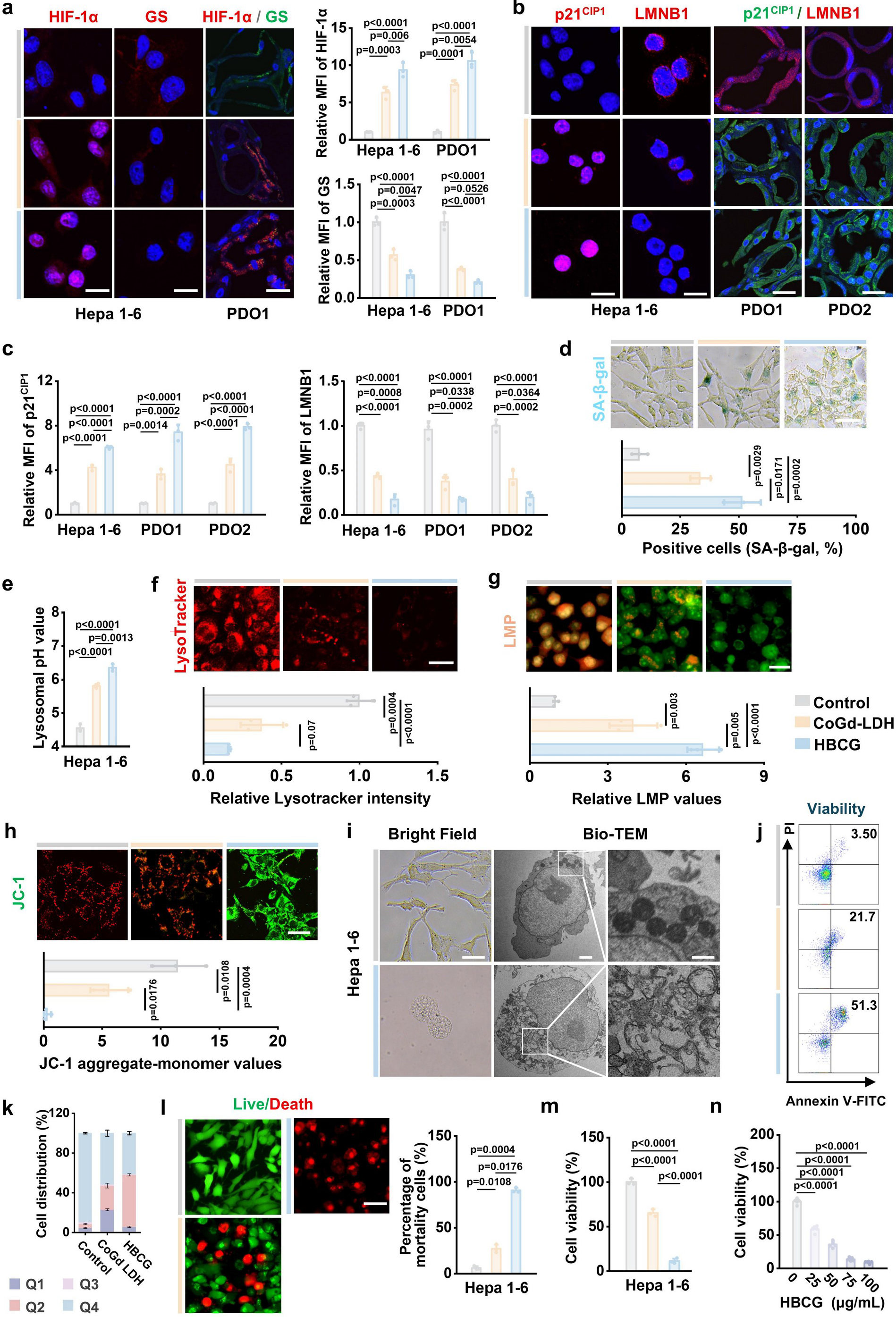
HBCG reinforces the ammonia–senescence ratchet and induces ammonioptosis in liver tumor models. **a**, Representative HIF-1α and GS immunofluorescence images and quantification in Hepa1-6 cells and PDO1 treated with control, CoGd-LDH or HBCG. **b**, **c**, Representative p21^CIP1^ and LMNB1 immunofluorescence images (b) and quantification (c) in Hepa1-6 cells, PDO1 and PDO2 after the indicated treatments. **d**, Representative SA-β-gal staining and quantification of SA-β-gal-positive Hepa 1-6 cells after the indicated treatments. **e**, Lysosomal pH in Hepa 1-6 cells after the indicated treatments. **f**, LysoTracker fluorescence images and quantification of fluorescence intensity. **g**, Fluorescence images and quantification of lysosomal membrane permeabilization in Hepa 1-6 cells after the indicated treatments. **h**, Representative JC-1 fluorescence images and quantification of the aggregate-to-monomer fluorescence ratio in Hepa 1-6 cells after the indicated treatments. **i**, Representative bright-field and Bio-TEM images of Hepa 1-6 cells after treatment with control or HBCG. Scale bars, 20 μm for bright-field images and 2 μm and 500 nm for TEM images. **j**, **k**, Representative flow-cytometric plots of Annexin V-FITC/PI staining (**j**) and quantification of the distribution of Hepa 1-6 cells among the four quadrants (**k**) after treatment with control, CoGd-LDH or HBCG. **l**, Representative live/dead staining images and corresponding quantification of dead Hepa 1-6 cells after the indicated treatments. **m**, Viability of Hepa 1-6 cells after treatment with control, CoGd-LDH or HBCG. **n**, Viability of Hepa 1-6 cells after treatment with 0-100 μg/mL HBCG. Nuclei in a and b were counterstained with DAPI. Scale bars, 20 μm (a, b, f–h and l) and 100 μm (d) respectively. Images are representative of three independent experiments. Data are mean ± s.d.; *n* = 3 biologically independent samples per group. One-way ANOVA followed by Tukey’s multiple-comparisons test was used in a, c–h and l–n. Exact P values are shown in the corresponding panels.

**Extended Data Fig. 5.**
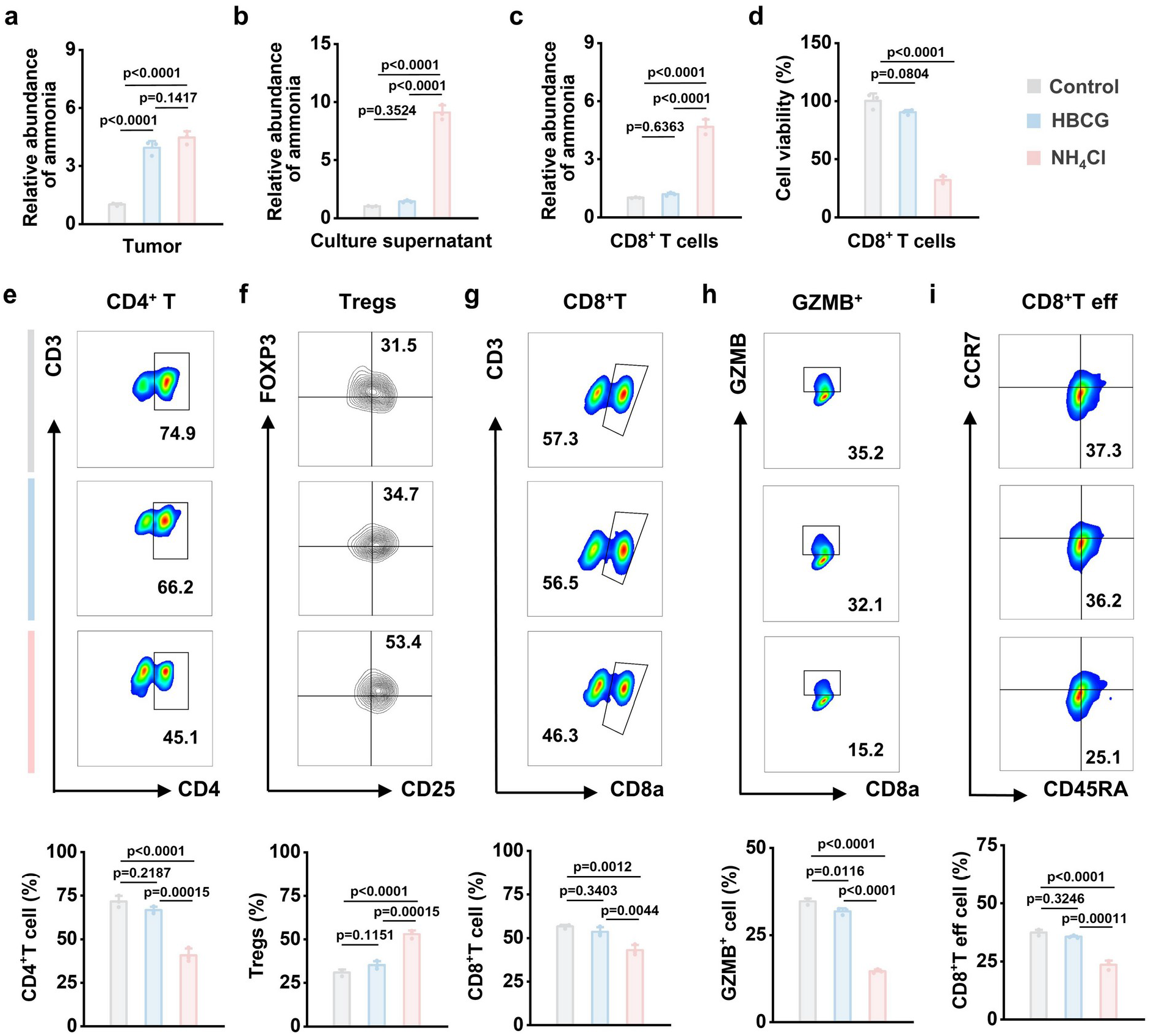
HBCG preferentially increases tumor-cell ammonia while limiting ammonia exposure and toxicity in human T cells. **a**–**c**, Relative ammonia abundance measured across different compartments of a tumor cell–PBMC co-culture system. Following treatment with control, HBCG or NH_4_Cl, tumor cells (**a**) and CD8⁺ T cells (**c**) were isolated by flow-cytometric sorting for intracellular ammonia measurement, while ammonia abundance in the shared culture supernatant was measured in parallel (**b**). **d**, Viability of CD8⁺ T cells following the indicated treatments. **e–i**, Representative flow-cytometry plots and corresponding quantification of CD3⁺CD4⁺ T cells (**e**), CD25⁺FOXP3⁺ Treg cells within the CD4⁺ T-cell compartment (**f**), CD3⁺CD8α⁺ T cells (**g**), GZMB⁺ cells within the CD8⁺ T-cell compartment (**h**) and CCR7/CD45RA-defined CD8⁺ effector T cells (**i**). Numbers in the representative flow-cytometry plots indicate the percentages of cells within the indicated gates. Data are mean ± s.d.; individual points represent independent samples, with *n* = 3 per condition. Statistical significance was assessed using one-way ANOVA followed by Tukey’s multiple-comparisons test. Exact *P* values are shown in the corresponding panels.

**Extended Data Fig. 6.**
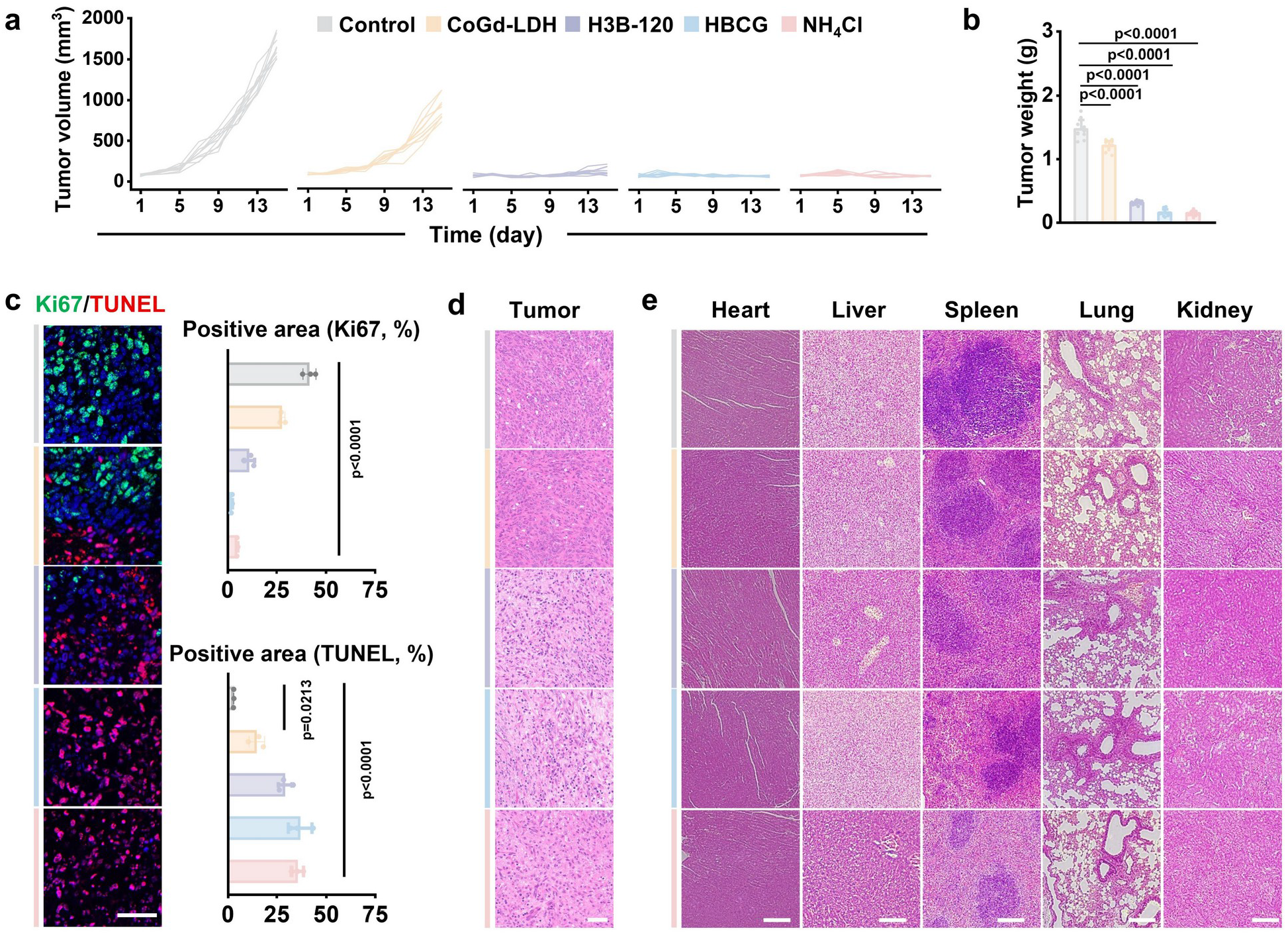
HBCG enforces the senescence–ammonioptosis axis and drives potent tumor elimination. **a**, Individual tumor growth curves of mice bearing subcutaneous tumors after intratumoral treatment with Control, CoGd-LDH, H3B-120, HBCG or NH_4_Cl on days 1, 3 and 5. Tumor volumes are shown for individual mice, n = 10 mice per group. **b**, Excised tumor weights at day 14 after the indicated treatments, n = 10 mice per group. **c**, Representative immunofluorescence images of Ki67 and TUNEL in tumor sections at day 14, with corresponding quantification of Ki67-positive and TUNEL-positive cells. Representative images are from three biologically independent mice per group. Quantification was performed using n = 3 biologically independent tumor samples per group. Scale bar, 50 μm. **d**, **e**, H&E staining of tumors (**d**) and major organs (heart, liver, spleen, lung, kidney) (**e**) after the indicated treatments. Scale bars, 100 μm. Data are presented as mean ± s.d. Statistical significance in b and c was determined by one-way ANOVA followed by Tukey’s multiple-comparisons test. All tests were two-sided unless otherwise indicated. Exact P values are provided in the panels.

